# Disentangling the genetic basis of rhizosphere microbiome assembly in tomato

**DOI:** 10.1101/2021.12.20.473370

**Authors:** Ben O Oyserman, Stalin Sarango Flores, Thom Griffioen, Xinya Pan, Elmar van der Wijk, Lotte Pronk, Wouter Lokhorst, Azkia Nurfikari, Nejc Stopnisek, Anne Kupczok, Viviane Cordovez, Víctor J Carrión, Wilco Ligterink, Basten L Snoek, Marnix H Medema, Jos M Raaijmakers

## Abstract

Microbiomes play a pivotal role in plant growth and health, but the genetic factors involved in microbiome assembly remain largely elusive. Here, 16S amplicon and metagenomic features of the rhizosphere microbiome were mapped as quantitative traits of a recombinant inbred line population of a cross between wild and domesticated tomato. Gene content analysis of prioritized tomato QTLs suggested a genetic basis for differential recruitment of various rhizobacterial lineages, including a *Streptomyces*-associated 6.31-Mbp region harboring tomato domestication sweeps and encoding, among others, the iron regulator FIT and the aquaporin SlTIP2.3. Within metagenome-assembled genomes of the rhizobacterial lineages *Streptomyces* and *Cellvibrio*, we identified microbial genes involved in metabolism of plant polysaccharides, iron, sulfur, trehalose, and vitamins, whose genetic variation associated with either modern or wild tomato QTLs. Integrating ‘microbiomics’ and quantitative plant genetics pinpointed putative plant and reciprocal microbial traits underlying microbiome assembly, thereby providing the first step towards plant-microbiome breeding programs.

## 1. Main

Root and shoot microbiomes are fundamental to plant growth and plant tolerance to (a)biotic stress factors. The outcome of these beneficial interactions is the emergence of specific microbiome-associated phenotypes (MAPs)^1^, such as drought resilience^2^, disease resistance^3^, development^4^ and heterosis (i.e. hybrid vigor)^5^. The microbes inhabiting the surface or internal tissues of plant roots are selectively nurtured by diverse plant-derived compounds in the form of primary and secondary metabolites^6,7^. Microbes reciprocate by supporting plant growth and producing metabolites that mediate processes such as nutrient acquisition and pathogen suppression^8,9^. Developing a blueprint of the genetic architecture for this ‘chemical dialogue’ and how these interactions lead to specific MAPs is a one of the key focal points in current plant microbiome research. The promise is that these genomic and chemical blueprints can be integrated into microbiome breeding programs for a new generation of crops that can rely, in part, on specific members of the microbiome for stress protection, enhanced growth and higher yields^10^.

Selective breeding for yield-related traits has left a considerable impact on the taxonomic and functional composition of modern crop microbiomes^11,12^. Wild plant relatives represent a ‘living library’ of diverse genetic traits that may have been lost during domestication^13^. For example, recombinant inbred lines (RILs) of crosses between wild tomato relatives and modern tomato cultivars have been used to identify genetic loci controlling important agronomic traits, including tolerance to abiotic^14^ and biotic stress^15^, as well as nutritional quality and flavor profiles^16^. To date, microbiome traits are not yet considered for breeding purposes, except for specific quantitative MAPs such as the number of nodules in legume-rhizobia symbioses^17^. However, technological advances in sequencing now make it feasible to treat microbiomes as quantitative traits for selection. This approach has been adopted for the phyllosphere microbiome and, recently, for the *Arabidopsis* and sorghum rhizosphere microbiomes^18,19^. For most plant species, however, investigations leveraging diverse plant populations to map microbiome Quantitative Trait Loci (QTL) are still at their infancy^20,19,18^. In these recent studies, the microbiomes were characterized by amplicon sequencing to detect loci involved in alpha and beta diversity as well as individual OTU abundances^21^. These studies provide strong evidence that microbiome recruitment has a genetic component, but the functional nature of the corresponding plant-microbe interactions cannot be elucidated from amplicon data. Hence, functional genomic features of the microbiome as well as intraspecific diversity within microbial species have not yet been taken into account^22^.

Here, we used both amplicon and shotgun metagenome sequencing to generate taxonomic as well as functional microbiome features as quantitative traits. Using an extensive recombinant inbred line (RIL) population of a cross between modern *Solanum lycopersicum* var. Moneymaker and wild *Solanum pimpinellifolium*^23^, we were able to identify reciprocal associations between specific plant and microbiome traits and to infer putative mechanisms for rhizosphere microbiome assembly. While both wild and modern alleles were identified, the large number of QTLs linked to modern alleles suggests that domestication has had a significant impact on rhizosphere microbiome assembly. The plant traits identified were related to growth, stress, amino acid metabolism, iron and water acquisition, hormonal responses, and terpene biosynthesis, whereas the microbial traits were related to metabolism of plant cell wall polysaccharides, vitamins, sulfur, and iron. Furthermore, we show that amplicon-based approaches allow detection of QTLs for rarer microbial taxa, whereas shotgun metagenomics allowed mapping to smaller and thus more defined plant genomic regions. Together, these results demonstrate the power of an integrated approach to disentangle and prioritize specific genomic regions and genes in both plants and microbes associated with microbiome assembly.

## 2. Results

### 2.1 Baseline analyses of the tomato Recombinant Inbred Line population

Prior to detailed metagenome analyses of the microbiome of the tomato RIL population, we first investigated whether QTLs previously identified in the same RIL population under sterile *in vitro* conditions could be replicated in our experiment conducted under greenhouse conditions with a commercial tomato greenhouse soil (Figure 1A and B, Supplemental table 1)^24^. We identified QTLs for Shoot Dry Weight (SDW) coinciding with a QTL identified previously on chr9^24^. Similarly, we identified QTLs for Rhizosphere Mass (RM), defined here as a the total mass of the roots with tightly adhering soil, which coincide with root trait QTLs previously identified for lateral root number, fresh and dry shoot weight, lateral root density per branched zone and total root size (Figure 1B)^24^. An analysis of variance (ANOVA) yielded significant variation in SDW based on the additivity of alleles linked to SDW (zero, one or two alleles) (F(2, 186) = 16.02, p = 3.76 e-07) (Figure 1C and 1D). A post hoc Tukey test further demonstrated significant differences between all pairwise comparisons (p < 0.05). For RM, an ANOVA yielded a significant difference (F(2, 186) = 16.02, p = 3.76 e-07); a post hoc Tukey test demonstrated a statistically significant difference only between presence of either one or two alleles (p < 0.05), but did not support additivity (p = 0.15) (Figure 1E and 1F). Collectively, our results confirm and extend earlier work conducted on the same tomato RIL population *in vitro*^24^, providing a solid basis for QTL mapping of taxonomic and genomic features of the rhizosphere microbiome.

**Figure 1:**
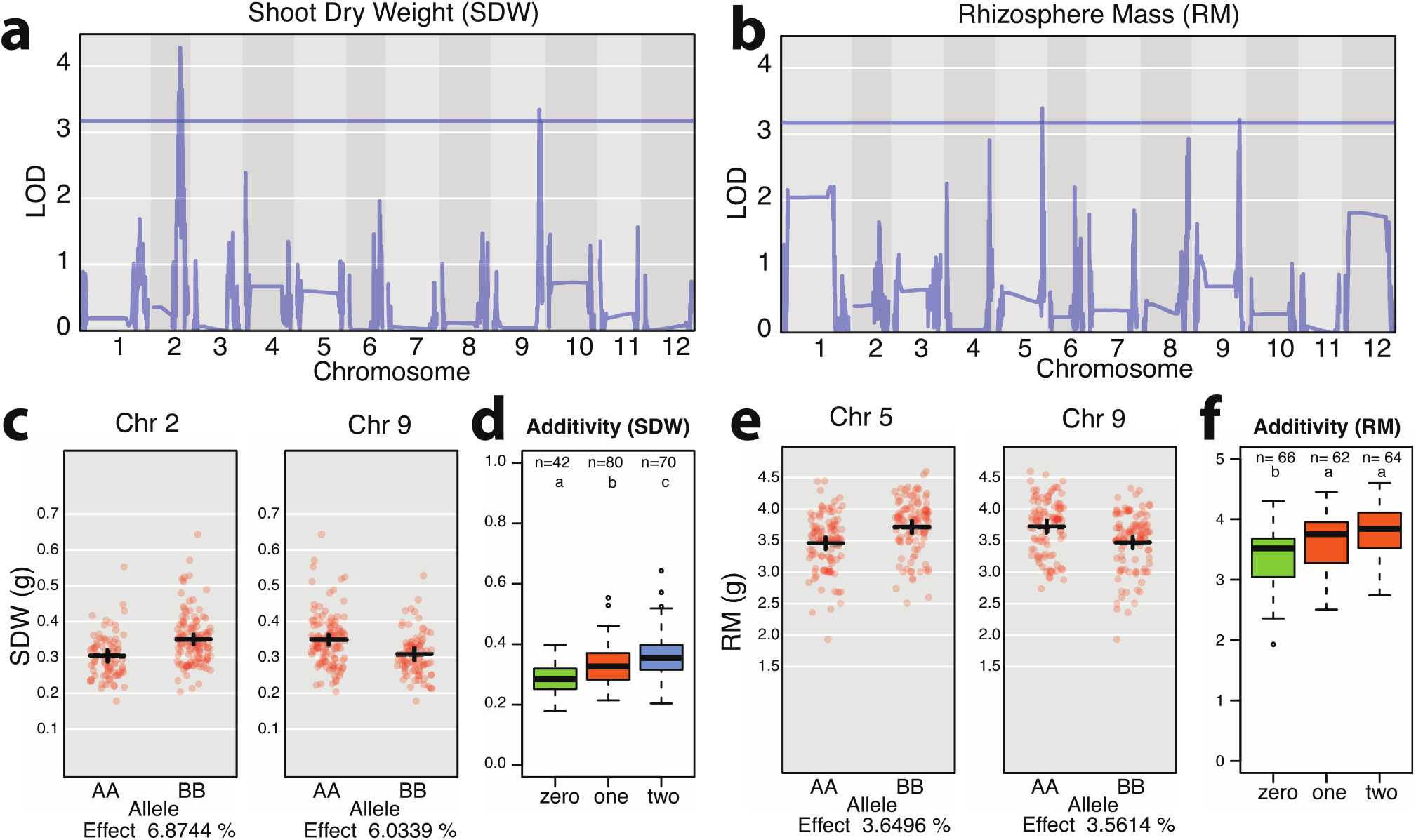
Identification of shoot dry weight (SDW) and rhizosphere mass (RM) QTLs in the recombinant inbred line (RIL) population of tomato. **(a)** QTLs identified for SDW on chromosome 9 position 63.63719184 and on chromosome 2 position 42.7291229, coinciding with a QTL identified previously (chromosome 9 position 62.897108) by Khan et al 2012. **(b)** QTL of RM on chromosome 5 position 62.00574891, and chromosome 9 position 62.71397636, which coincide with root trait QTLs previously identified by Khan et al 2012 for lateral root number chromosome 5 position 53.4-86.1, and several on chromosome 9, including fresh and dry shoot weight, (chromosome 9 position 81.3-95.3), lateral root density per branched zone (chromosome 9 position 33.8-88.7), and total root size (chromosome 9 position 39.4-75.1). **(c)** Scatter plots showing the distribution of SDW measurements on chromosome 2 position 42.7291229 and chromosome 9 position 63.63719184 for both modern (AA) and wild (BB) tomato alleles. **(d)** Significant additivity of tomato alleles for shoot dry weight (p < 0.05); n of 42, 80 and 70 for tomato plants containing neither allele (labeled zero), either BB allele on chromosome 2, or AA on chromosome 9 (labeled one), or both AA and BB alleles (labeled two), respectively. **(e)** Scatter plots showing the distribution of RM measurements on chromosome 5 (pos 62.00574891), and chromosome 9 (pos 62.71397636) for both modern (AA) and wild (BB) alleles. **(f)** No additivity of alleles was observed for RM.

### 2.2 Taxonomic microbiome features as quantitative traits

To investigate molecular features of the microbiome as quantitative traits, we conducted 16S amplicon sequencing of 225 rhizosphere samples, including unplanted bulk soil, parental tomato genotypes, and all 96 RIL accessions in duplicate (Supplemental table 2-5, BioProject ID PRJNA787039). We observed a separation between the microbiomes of rhizosphere and bulk soil, between the microbiomes of the two parental tomato genotypes and the RIL accession microbiomes (Figure 2A). To limit multiple testing and to focus on common microbiome features with sufficient coverage across all accessions, we prioritized the rhizosphere-enriched amplicon sequence variants (ASVs) to those present in 50% or more of the RIL accessions (Figure 2B). A QTL analysis with these prioritized ASVs was run with R/qtl2^25^ using a high-density tomato genotype map^26^, harvest date, post-harvest total bulk soil mass, RM, number of leaves at harvest and SDW as co-variates.

**Figure 2:**
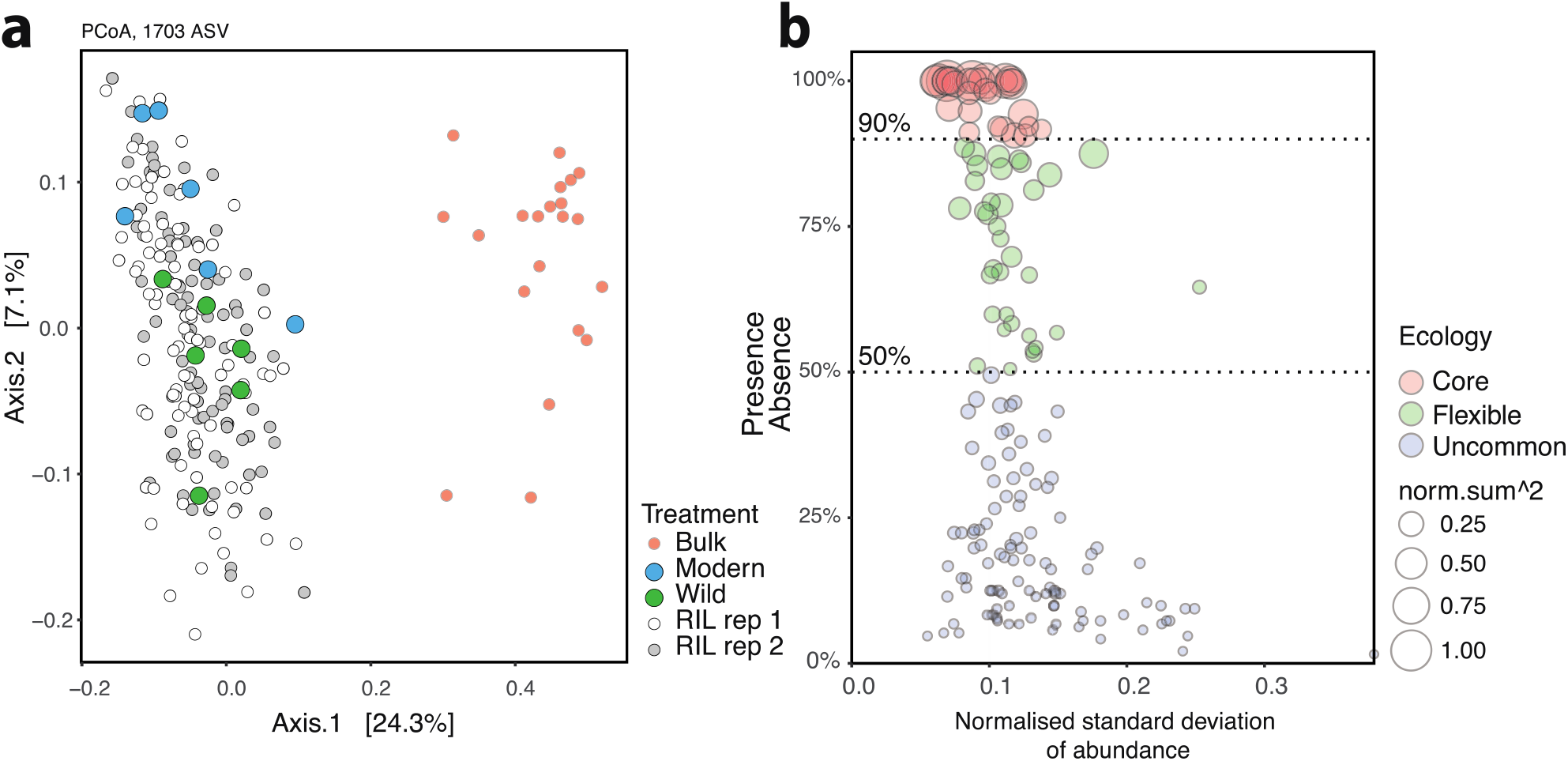
PCoA analysis of the 16S rRNA amplicon data obtained for the microbiomes of bulk soil and the rhizosphere of modern and wild tomatoes and their recombinant inbred line (RIL) population. **(a)** PCoA analysis of amplicon sequence variants (ASVs) demonstrating a separation between the bulk soil and rhizosphere microbiomes. The rhizosphere microbiome of the 96 RIL accessions distributed around those of the wild and modern rhizosphere microbiomes. Separation between the two replicate RIL populations was not observed. **(b)** To limit multiple testing, a QTL analysis was conducted only on ASVs that were observed in more than 50% of the RIL accessions.

We identified 48 QTL peaks, across 45 distinct loci, significantly associated with 33 ASVs (Supplemental table 6). Our logarithm of the odds (LOD) thresholds for significance had been determined by pooled permutations from all ASVs to attain a genome-wide threshold of P 0.05 (LOD 3.35) and P 0.2 (LOD 2.64). Of the significant QTLs, 16 were more abundant in a wild tomato allele and 32 in a modern tomatos allele. The QTLs on chromosomes 11, 10, 8 and 2 were all linked to ‘modern’ alleles; the sole QTL on chromosome 7 was linked to a ‘wild’ tomato allele. All other chromosomes contained a mix of QTLs linked to either modern or wild alleles (Figure 3A). While many rhizobacterial lineages were linked to a single QTL (14 taxa out of 25), others were linked to two or more QTLs (7 and 4 taxa, respectively) (Figure 3B). Of the lineages with multiple QTLs, most were linked only to modern tomato alleles. One salient exception was *Methylophilaceae*, with increased abundance linked to a total of 9 QTLs, from both wild and modern alleles, and distributed across chromosomes 3 (modern, x2), 4 (modern), 7 (wild), 11 (modern x2) and 12 (wild x3) (Figure 3D). Another salient feature of the QTL analysis was the hotspot for microbiome assembly identified on chromosome 11, including ASVs from *Adhaeribacter, Caulobacter, Devosia*, Rhizobiaceae, *Massilia* and *Methylophilaceae* (Figure 3D).

**Figure 3:**
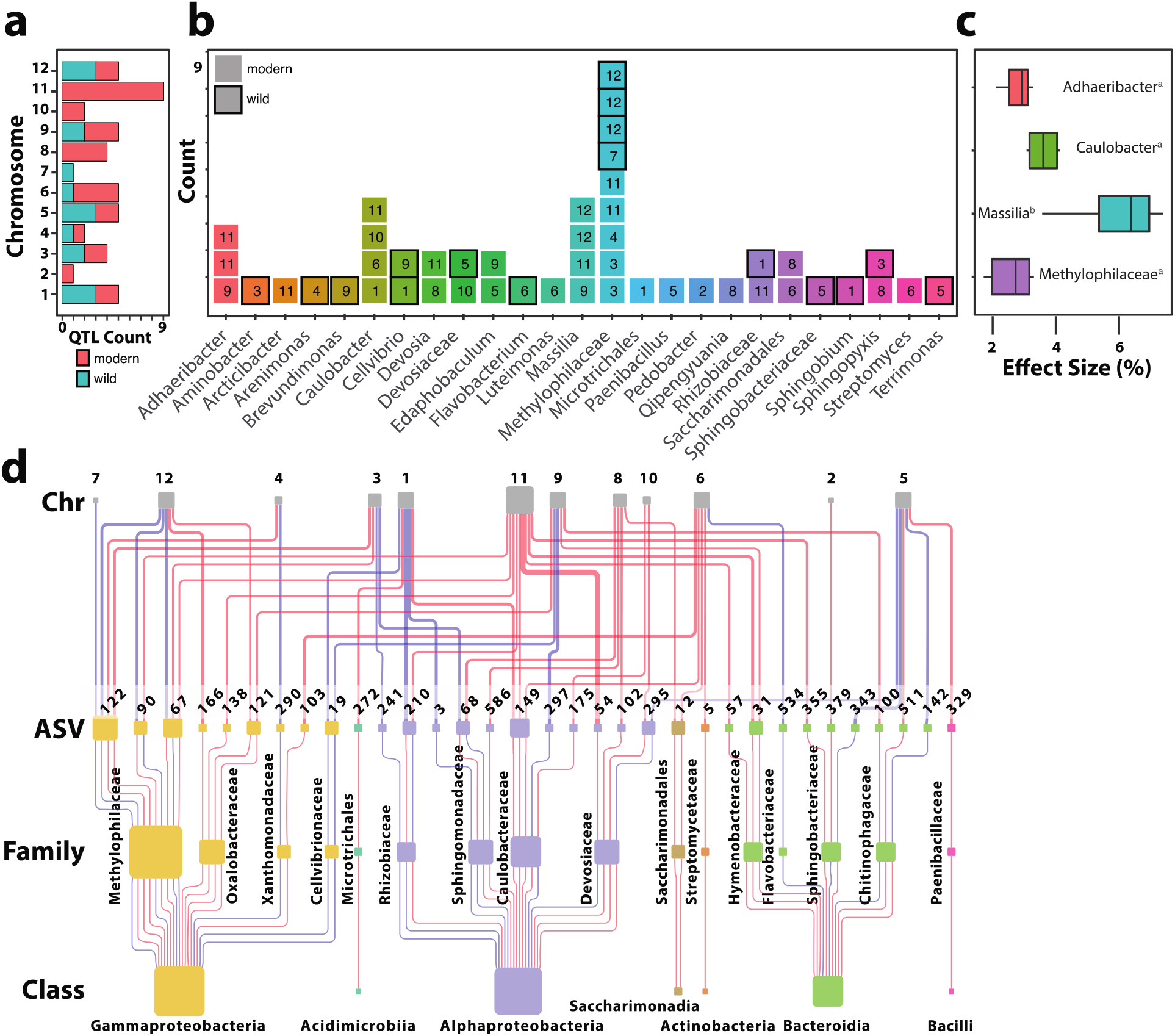
Association between 16S rRNA amplicon sequence variants (ASVs) and tomato QTLs **(a)** A color-coded summary of the number of 16S rRNA QTLs identified per chromosome of wild and modern tomato alleles. **(b)** A summary of the number of 16S rRNA QTLs linked to bacterial taxonomies, with the chromosome number of each QTL represented within each square. The presence and absence of dark borders around each square are used to indicate a QTL linked to higher abundance for a wild allele and modern allele, respectively. **(c)** Effect size for four rhizobacterial lineages with 3 or more QTLs. **(d)** Hierarchical network depicting the 16S rRNA QTLs identified in this study. From top to bottom: the first row represents tomato chromosomes (Chr), which are linked to specific ASVs in the next row, which taxonomically belong to different families and classes of bacteria in subsequent rows. The size of the chromosome nodes is weighted by the number of outbound edges. The ASV, family, and class node sizes are weighted by the number of in-bound edges. A complex network emerges, whereby the abundance of individual ASVs, at different taxonomic levels, is determined by a network of interactions of multiple tomato alleles from both modern and wild origin.

The effect size of the 48 QTLs on ASV relative abundance ranged from 1.3 to 17%, with an average effect size of approximately 5%, comparable to the effects seen for SDW and RM (Figure 1C and E). The largest effect was a single modern QTL for an ASV in the genus *Qipengyuania* (17%), and a second modern QTL in *Edaphobaculum* (10%). No statistical difference was found between modern and wild alleles on their effect size (p = 0.78, two-tailed t-test). For those lineages with sufficient representation at the class level (Bacteroidia, Alphaproteobacteria, and Gammaproteobacteria), there was no statistically significant difference between effect size (F(3, 16) = 0.072, p = 0.974). However, an ANOVA on the positive effect size at genus level demonstrated significant differences between lineages (F(3, 16) = 12.94, p = 1.15 e-04). A post hoc Tukey test demonstrated QTLs for *Massilia* with a larger positive effect size than other lineages with sufficient sample size for comparison (Figure 3C). Together, the amplicon analysis provided a broad picture, suggesting that microbiome assembly is a complex trait governed by a combination of multiple loci, some being ASV specific, some being pleiotropic to different ASVs and with heterogenous effect sizes (Figure 3D). While positive effects were identified linked to both wild and modern alleles, the large number of QTLs linked to modern alleles, suggests domestication has had a significant impact on rhizosphere microbiome assembly.

### 2.3 Functional microbiome features as quantitative traits

To understand the functional traits associated with rhizosphere microbiome assembly, we generated shotgun metagenomes for each accession in the tomato RIL population (96 total), as well as six samples of the modern tomato parent, five samples of the wild tomato parent and seven bulk soil samples (BioProject ID PRJNA789467). After pre-processing, assembly, back-mapping, CSS normalization and binning, QTL mapping was conducted for the rhizosphere enriched contig and bin abundances. Binning was done using Metabat2 (version 2:2.15)^27^ and genomic quality of the output was evaluated by CheckM^28^ (Supplemental Table 7). The bins and assembled contigs larger than 10kb can be found on Open Science Framework (https://osf.io/f45ek/). All contigs of 10kb and larger were taxonomically assigned using Kraken^29^ (Supplemental Table 8). With nearly 40 million contigs being assembled, we took numerous prioritization steps to reduce the effects of multiple testing. Only rhizosphere-enriched contigs larger than 10kb and with a rhizosphere enrichment greater than 4-fold were selected resulting in 1249 contigs. Only bins with greater than 90% completion and less than 5% contamination were mapped (33 out of 588 bins). As with the ASVs, harvest date, bulk soil mass, rhizosphere mass (RM), number of leaves at harvest, and SDW were used as co-variates in QTL mapping (supplemental table 11 and 12, respectively).

We identified 7 significant bin QTLs (LOD > 3.40, P < 0.05) (Supplemental table 9) including *Streptomyces* bin 72 associated with a modern allele on tomato chromosomes 6 and 11. For the contigs, a total of 717 QTLs at 26 unique positions on chromosomes 1, 4, 5, 6, 9 and 11 were identified (Supplemental table 10), corresponding to 476 metagenomic contigs from 10 different genera (LOD > 3.47, P < 0.05). The largest number of contig QTLs belonged to the *Streptomyces, Cellvibrio* and *Sphingopyxis* lineages (Figure 4A). The *Streptomyces* contigs mapped to QTLs on tomato chromosomes 4 (46 contigs, wild tomato), 6 (190 contigs, modern tomato) and 11 (257 contigs, modern tomato), with a subset of contigs mapping to two or all three of these positions (Figure 4B). These findings corroborate and expand upon the *Streptomyces* QTL identified on chromosome 6 using our 16S amplicon data, as well as that of the bin QTLs identified on chromosomes 6 and 11. The *Cellvibrio* contigs mapped to chromosome 1 (42 contigs, wild) and chromosome 9 (94 contigs, wild), again corroborating the findings from our 16S amplicon analysis described above. In contrast, the *Sphingopyxis* QTLs identified on chromosome 5 (24 contigs, wild) and 9 (49 contigs, modern) did not correspond to the QTLs identified on chromosomes 8 and 3 in the 16S amplicon analysis. Interestingly, 4 contigs for *Devosia* also corroborated the results of the 16S QTL analysis. The effect sizes ranged from 9 to 21 % and were significantly different (F(14, 702) = 530.9 p < 2e-16) between QTL and lineages (Figure 4C). Interestingly, as with the 16S amplicon analysis, some of the highest LOD scores were for contigs belonging to *Devosia*. Also, the effect size of the *Sphingopyxis* contigs was large (± 20% on average), above 15% for *Cellvibrio*, and approximately 10% for *Streptomyces*. The average QTL region was 51.59 Mbps for the 16S amplicon sequences and 26.64 Mbps for the metagenomic contigs (two-sided t.test, p = 3.32E-09) (Figure 4E). A more striking contrast was observed in the difference between the median size of amplicon and contig QTL regions which were 58.56 Mbp and only 6.47 Mbp, respectively. In summary, while many more taxa were identified in the amplicon-based QTL analysis, the metagenome-based QTL analysis provided QTLs with much smaller confidence intervals (Figure 4E).

**Figure 4:**
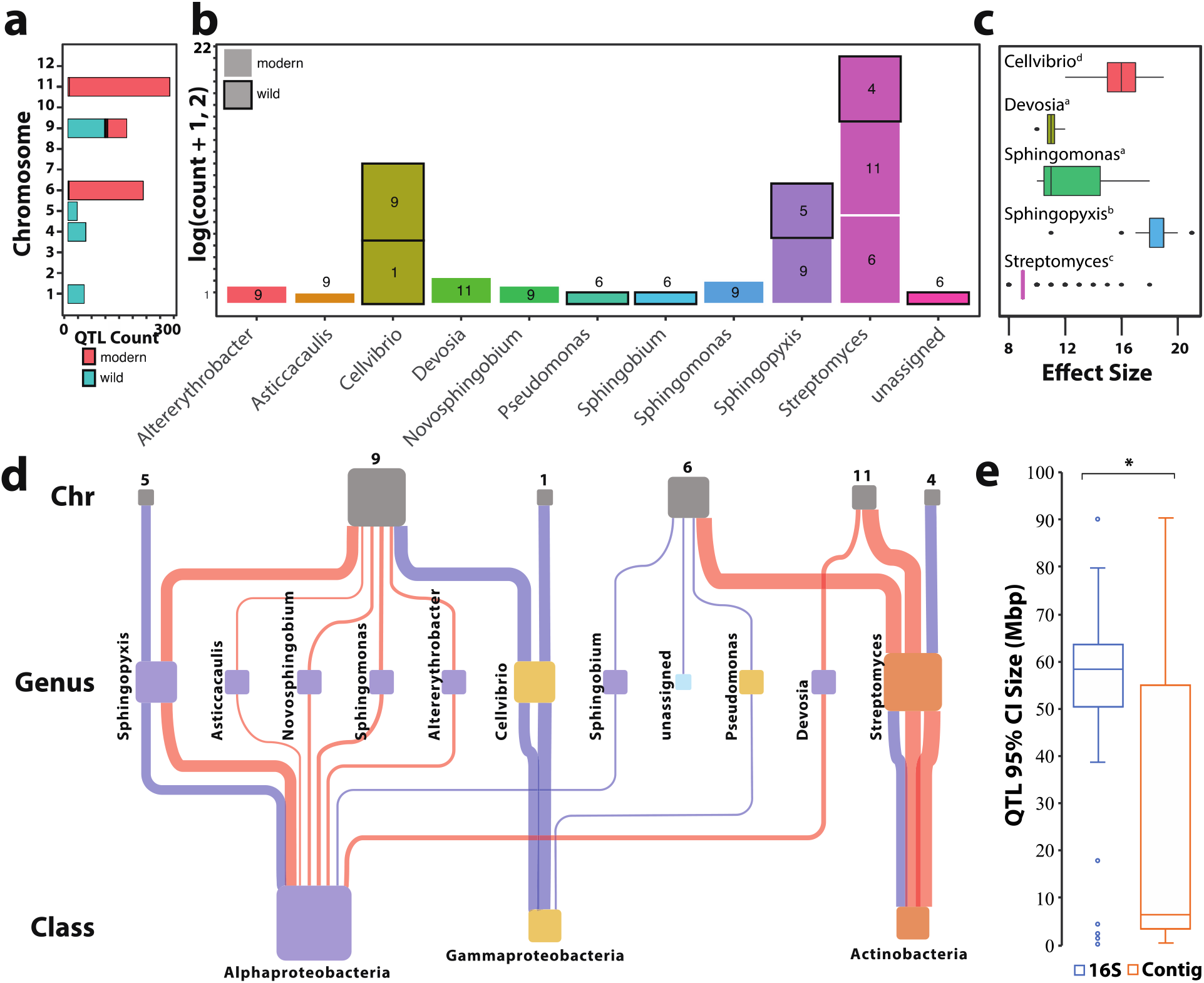
Association between metagenomic contigs of the rhizosphere microbiome and tomato QTLs **(a)** A color coded summary of the number of contig QTLs identified per chromosome to wild and modern alleles. **(b)** A summary of the number of contig QTLs found by taxonomies, with the chromosome of each QTL represented within each square. The presence and absence of dark borders around each square are used to indicate a QTL linked to higher abundance for a wild allele and modern allele, respectively. **(c)** The effect sizes for each lineage were significantly different as indicated by letters (F(14, 702) = 530.9 p < 2e-16) **(d)** A hierarchically structured network depicting the contig QTLs identified in this study. From top to bottom are the tomato chromosomes (Chr), which are associated with specific metagenomic contigs and taxonomically linked to different families and classes of bacteria. The size of the chromosome nodes is weighted by the number of outbound edges. The ASV, family, and class node sizes are weighted by the number of in-bound edges. **(e)** Comparison between the size of the QTL regions identified based on 16S amplicon data and based on metageonomic contigs. The 95% confidence interval of contig QTLs was significantly smaller than the 95% confidence interval of 16S rRNA QTLs (two-sided t.test, p = 3.32E-09).

### 2.4 Amplicon-based bulk segregant analysis of Streptomyces and Cellvibrio abundance

The two most abundant rhizosphere taxa with replicated patterns for amplicon and metagenome-based QTLs were *Streptomyces* and *Cellvibrio*. Therefore, we sought to provide additional independent support for these QTLs using a bulk segregant analysis of an independent population of parental and RIL genotypes (Supplemental_Table_11). In particular, we tested the previously identified amplicon-based QTLs associated with higher *Cellvibrio* abundance at markers 464 and 3142 on chromosomes 1 and 9, respectively with higher *Streptomyces* abundance at marker 2274 on chromosome 6 (Figure 5). In each case, ANOVA showed a statistical difference between genotypes and bulk soil, respectively (F(4, 396) = 21.56, p = 4.16 e-16), (F(4, 396) = 18.43, p = 6.68 e-14), (F(4, 396) = 8.423, p = 1.57 e-06). A post hoc Tukey test supported the conclusion that wild allele at markers 464 and 3142 on chromosomes 1 and 9, respectively, are indeed associated with increased abundance *Cellvibrio* (p = 3.913 e-04, and p = 0.08 respectively), while the modern allele at markers 2274 on chromosome 6 was significantly associated with increased abundance of *Streptomyces* (p = 1.152 e-04).

**Figure 5:**
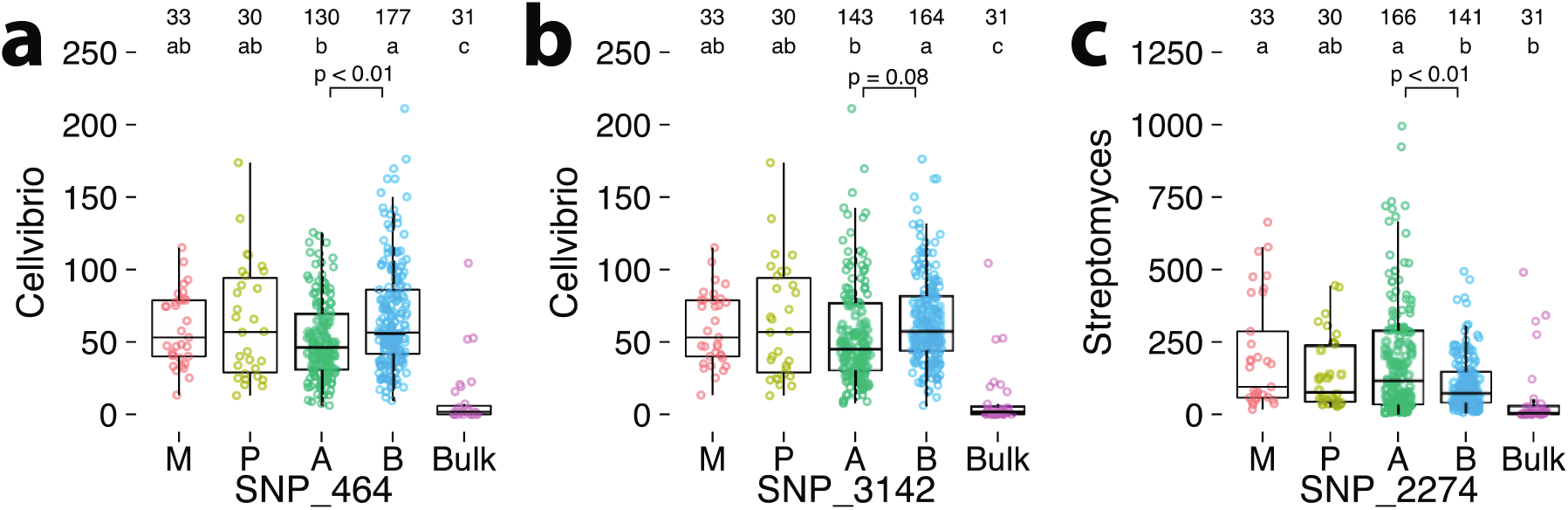
Validation of Cellvibrio and Streptomyces 16S rRNA QTLs with bulk segregant analysis in an independent experiment with modern, wild and 77 RIL accessions (see Supplemental table 13). The number of replicates for each treatment is detailed in the top row of each panel. The number of replicates within the RIL population are represented by either an A (modern) or B (wild) allele, which depends on the marker in question. The row below represents the statistical group based on Tukey’s HSD; a different letter indicates a statistically significant difference. **(a)** The relative abundances of Cellvibrio 16S rRNA in bulk soil, modern, wild, and RIL accessions at SNP marker position 464 on chromosome 1. At this position, 32 and 45 RIL accessions with modern and wild alleles were used (130 and 177 samples with biological replication respectively). **(b)** Similarly, for SNP marker 3142 on chromosome 9, there were a total of 35 and 42 RIL accessions with modern and wild alleles, (143 and 164 samples with biological replication respectively). **(c)** The relative abundances of Streptomyces 16S rRNA and sequences in bulk soil, modern, wild, and RIL accessions at SNP marker 2274 on chromosome 6. There was a total of 42 and 35 RIL accessions with modern and wild alleles, (166 and 141 samples with biological replication, respectively).

### 2.5 Host genetics and rhizosphere microbiome assembly

A subset of 5 regions consistent across both the amplicon and metagenome-based analyses were prioritized with an average size of 2.68 Mbps (Supplemental Table 12). These included positions on chromosome 1 (positions 87.36 - 90.49 Mbps), chromosome 9 (pos 62.03 – 63.32 Mbps), chromosome 5 (pos 61.54 – 63.38), chromosome 6 (pos 33.99 – 40.3 Mbps) and chromosome 11 (pos 53.06 - 53.89 Mbps). In total, 1359 genes were identified in these regions. Potential candidate genes with root-specific transcriptional patterns, defined as a 4 fold increase in the roots compared to leaf samples, were further prioritized using a publicly available RNAseq dataset^30^. Based on this analysis, a subset of 192 root specific genes were identified (Supplemental table 13). A total of 98 root specific genes were linked to *Streptomyces* on chromosome 6 (84 genes) and 11 (14 genes) (Figure 6). Intriguingly, 61 of these genes were found in regions previously identified to be subjected to selective sweeps related to tomato domestication as well as to subsequent sweeps related to improvements in fruit quality^31^(Supplemental Figure 1).

**Figure 6:**
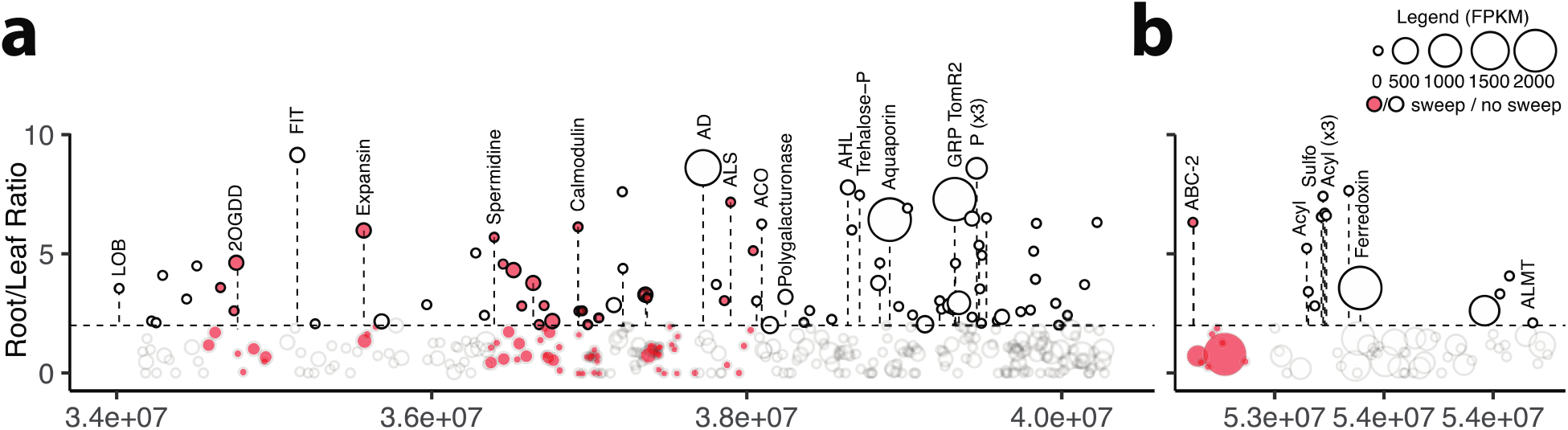
Prioritized regions of the Streptomyces-associated QTLs on tomato chromosomes 6 and 11 overlaying previously published data^30^ on root-specific gene expression and genetic sweeps due to domestication^31^ (in red). Within each region, the log_2_ ratio gene expression patterns from leaf and root materials were calculated and those with a log_2_ greater than 2, as delineated by the dotted line, were further prioritized. The log_2_ root transcript abundances are depicted by the size of the bubble. **a)** The 6.31 Mbp region on chromosome 6, position 33.99-40.3 Mbps. Abbreviations of highlighted genes: LOB - LOB domain protein 4, 2OGDD - 2-Oxoglutarate-dependent dioxygenases, FIT - FIT (Fer-like iron deficiency-induced transcription factor), Spermidine - Spermidine synthase, AD - Alcohol dehydrogenase 2, ALS - Acetolactate synthase, ACO - 1-aminocyclopropane-1-carboxylate oxidase, Polygalacturonase, AHL - AT-hook motif nuclear-localized protein, Trehalose-P - Trehalose 6-phosphate phosphatase, Aquaporin - Tonoplast intrinsic protein 23 / Aquaporin, GPR TomR2 - Glycine-rich protein TomR2, P - Acid phosphatase (x3). **b)** The 0.83 Mbp region on chromosome 11, position 53.06-54.89 Mbps. Abbreviations of highlighted genes: ABC-2 - ABC-2 type transporter, Acyl – Acyltransferase (x4), Sulfo – Sulfotransferase, ALMT-Aluminum-activated malate transporter.

Two of the most salient genes in this list included genes with high transcription in the roots; an aquaporin and a Fer-like iron deficiency-induced transcription factor (FIT). The aquaporin (SlTIP2.3) is one of eleven tonoplast intrinsic proteins in the tomato genome^32^ and has the highest fold change in the roots compared to all other organs^33^. The FIT gene is a bHLH transcriptional regulator controlling iron homeostasis in tomato^34,35^. Other genes of interest on chromosome 6 include a glycine rich protein, a receptor like kinase known to be upregulated during drought^36^, alcohol dehydrogenase, numerous phosphatases, expansins, ethylene-responsive transcription factors, gibberellin receptors, aminocyclopropane-1-carboxylate oxidase (ACO), an enzyme involved in the last step of ethylene biosynthesis, and finally, alpha-humulene and (-)-(E)-beta-caryophyllene, a known tomato terpene and signaling molecule in tomato^37,38^ and also acting as a volatile in microbiome assembly^39^. Root specific genes involved in carbohydrate, protein and amino metabolism were also identified, including trypsin-alpha amylase inhibitor, prolyl 4-hydroxylase, polygalacturonase, trehalose phosphatase, glycogenin, xyloglucan fucosyltransferase and a metallocarboxypeptidase inhibitor, spermidine synthase, acetolactate synthases, alanine aminotransferase, and an amino acid permease. On chromosome 11, a ferrodoxin, an aluminum activated malate transporter^40^ and a cluster of various acetyltransferases and a sulfotransferase were identified.

A total of 57 root specific genes were identified in the QTL regions on chromosome 1 and 9 linked to *Cellvibrio*. These include a cytochrome p450 involved in coumarin synthesis, numerous extensins, phosphatases, respiratory burst oxidase-like protein, iron chelator nicotianamine synthase^41,42^ and on chromosome 11 phenazine biosynthesis. On chromosome 5, 37 root specific genes were identified including multiple peroxidases, glutamine synthetase, rhamnogalacturonate lyase, pectinesterase, metacaspase and trehalose-phosphatase. Furthermore, numerous ethylene responsive transcription factors and receptor like kinases were observed. The QTL on chromosome 1 contains genome-wide sweeps related to the initial tomato domestication and subsequent improvements of fruit quality traits, suggesting that one or both of these events was linked to a decreased abundance of root-associated *Cellvibrio*.

### 2.6 Illuminating metagenomic traits in Cellvibrio and Streptomyces

To further investigate the potential functional importance of the 476 rhizosphere-enriched metagenomic contigs mapped as QTLs, we performed a deeper analysis into their functional gene content (Supplemental Table 14-16). An antiSMASH^43^ analysis identified 30 biosynthetic gene clusters (BGCs) across these contigs. These BGCs largely originated from contigs taxonomically assigned to *Cellvibrio* and *Streptomyces*. They included several gene clusters potentially associated with root colonization, such as two melanin BGCs (c00216, NODE_5919; c00255, NODE_7250) from *Streptomyces* (which have been positively associated with colonization^44^) and a *Cellvibrio* aryl polyene BGC (c00185, NODE_4941), which is thought to protect bacteria against reactive oxygen species generated during immune responses of the host plant^45^. The contigs also contained gene clusters potentially beneficial to the host, such as BGCs encoding iron-scavenging siderophores, which have been associated with disease suppression in tomato^46^; specifically, homologues of coelichelin and desferrioxamine BGCs from streptomycetes were found (c00269, NODE_7969 and c00122, NODE_3362), three IucA/IucC-like putative siderophore synthetase gene clusters (c00106, NODE_2973; c00041, NODE_1131; c00238, NODE_6661), as well as a *Cellvibrio* NRPS-PKS gene cluster (c00001, NODE_101) most likely encoding the production of a siderophore based on the presence of a TonB-dependent siderophore receptor-encoding gene as well as a putative *tauD*-like siderophore amino acid β-hydroxylase-encoding gene^47^. The *Cellvibrio* contigs also contain several genes relevant for carbohydrate catabolism. For example, homologs of *xyl31a* (B2R_23365) and *bgl35a* (B2R_06825-06826) were detected (with 78%, 79% and 65% amino acid identity, respectively), genes that have been shown to be responsible for utilization of the abundant plant cell wall polysaccharide xyloglucan in *Cellvibrio japonicus*^48^. In addition, a possible homologue of the β-glucosidase gene *bgl3D*^49^ (B2R_26663), involved in xyloglucan utilization, was also identified, having high similarity to *bgl3D* from *Cellvibrio japonicus* (64% amino acid identity). Also, putative cellulose-hydrolizing enzymes were detected, such as a homologue (B2R_21082) of the cellobiohydrolase *cel6A* from *Cellvibrio japonicus*^50^ encoded in a complex locus of nine carbohydrate-acting enzymes annotated on this contig (NODE_5090) by DBCAN^51^ (Supplemental Table 14). Collectively, these results point to a possible role of microbial traits related to iron acquisition and metabolism of plant polysaccharides in tomato rhizosphere microbiome assembly.

Contigs of the metagenome-assembled genome (MAG) associated with *Streptomyces* ASV5 (the key taxon associated with tomato QTLs described above) contained a multitude of functional genes potentially relevant for host-microbe interactions. Taxonomically, the ASV5 MAG was most closely related to a clade of streptomycetes that includes type strains of species such as *arenae, flavovariabilis, variegatus*, and *chartreusis*. To understand how tomato might differentially recruit ASV5 streptomycetes, we analyzed the MAG for genes and gene clusters potentially involved in colonization. Intriguingly, we found contigs to be rich in genes associated with plant cell wall degradation. In particular, we identified a family 6 glycosyl hydrolases (B2R_10154) of which the glycosyl hydrolase domain has 84% amino acid identity to that of the SACTE_0237 protein that was recently shown to be essential for the high cellulolytic activity of *Streptomyces* sp. SirexAA-E^31^. Additionally, we detected a homologue (82% amino acid identity) of *Streptomyces reticuli* avicelase, a well-studied cellulase enzyme that degrades cellulose into cellobiose^52^ (B2R_29198). Larger gene clusters associated with degradation of plant cell wall materials were also found. These included an 8-kb gene cluster coding for multiple pectate lyases and pectinesterases (B2R_31553-31558), and an 8-kb gene cluster encoding a family 43 glycosyl hydrolase, a pectate lyase L, a rhamnogalacturonan acetylesterase RhgT, a GDSL-like lipase/acylhydrolase, a family 53 glycosyl hydrolase, and an endoglucanase A (B2R_15915-15920). Together, these findings suggest that ASV5 *Streptomyces* has the capacity to effectively process complex organic materials shed by plant roots during growth. These results are in line with a recent study on plant-associated streptomycetes that indicated that their colonization success appears to be associated with the ability to utilize complex organic material of plant roots^53^.

Root exudates also play a key role in recruitment of microbes. Prominent sugar components of tomato root exudates are glucose, but also xylose and fructose^54^. The *Streptomyces* MAG contains *xylA* and *xylB* genes (B2R_19014, B2R_19013) and a putative *xylFGH* import system (B2R_29274, B2R_23438, B2R_23439) facilitating xylose catabolization. Similarly, a *frcBCA* import system was identified in the genome (B2R_17966-B2R_17968) as well as a glucose permease (B2R_32780) with 91,5% amino acid identity to g*lcP1* SCO5578 of *Streptomyces coelicolor* A3(2)^55^. Other genes putatively involved in root exudate catabolism were also found in the ASV5 MAG, such as sarcosine oxidase (*soxBAG*, B2R_20550-20551 and B2R_21105), which has been shown to be upregulated in the presence of root exudates of various plants^56,57^.

In summary, the *Cellvibrio* and *Streptomyces* contigs encoded a range of functions that likely allow them to profit from tomato root exudates as well as complex organic material shed from growing tomato roots. How these plant traits differ between wild and domesticated tomatoes and if/how these influence differential colonization of roots of wild and domesticated tomato lines by these two bacterial lineages will require detailed comparative metabolomic analyses of the root exudates of both tomato lines as well as isolation of the corresponding *Cellvibrio* and *Streptomyces* ASVs, analysis of their substrate utilization spectrum followed by site-directed mutagenesis of the candidate genes, root colonization assays and *in situ* localization studies.

### 2.7 Genomic structure in Cellvibrio and Streptomyces provides insights into adaptations for differential recruitment

Bacterial populations often contain significant genomic heterogeneity. This heterogeneity may be associated with differential recruitment through altered nutrient preferences or host colonization mechanisms. The use of metagenomics enabled us to investigate the population structure within each rhizobacterial lineage and identify intraspecific differences. To do so, we first identified a unique set of 697,731 microbiome SNPs in a subset of parental and bulk metagenomes using InStrain^22^. A set of 15,026 SNPs enriched in either the wild or modern tomato rhizosphere were selected and the abundance of each allele at each SNP was calculated. Using these abundances, QTL mapping was performed using R/qtl2 as described in the methods. A total of 3,357 QTL peaks were identified (LOD > 3.01, P < 0.05), to 1229 independent loci. A total of 1,354 QTL peaks were more abundantly associated to a modern, and 2,001 to a wild plant allele, derived from 2,898 unique SNPs, and corresponding to 810 and 1,068 unique rhizobacterial genes respectively (Supplemental Table 17).

We investigated the 103 *Streptomyces* SNP QTLs at 94 unique positions within annotated genes whose mapping coincided with the previously identified *Streptomyces* contig QTLs to chromosomes 4, 6 and 11 (Supplemental Table 17). Numerous SNPs were associated with a higher abundance to the modern tomato alleles on chromosome 6 and 11. In particular, alpha-galactosidase (B2R_16136) and arabinose import (B2R_29105) had the highest LOD and smallest overlapping confidence intervals with chromosomes 6 and 11 (Figure 7). Indeed, many SNPs in genes involved in the degradation of xylan^58^, one of the most dominant non-cellulosic polysaccharides in plant cell-walls^59^, as well as carbohydrate and protein metabolism were linked to chromosomes 6 and 11, including xyloglucanase Xgh74A (B2R_10589), alpha-xylosidase (B2R_23763), endo-1,4-beta-xylanase (B2R_20609), extracellular exo-alpha-L-arabinofuranosidase (B2R_20608), multiple protease HtpX (B2R_19218), cutinase (B2R_19356), and putative ABC transporter substrate-binding protein YesO (B2R_09821) which has been implicated in the transport of plant cell wall pectin-derived oligosaccharides^60^. A *Streptomyces* SNP in acetolactate synthase (B2R_28001) was linked to chromosome 6 where a plant acetolactate synthase was located. Similarly, multiple SNPs in *Streptomyces* genes involved in putrescine transportation (B2R_25489) were linked to chromosomes 6 and 11, which contain genes for spermine synthase, suggesting a possible metabolic cross-feeding from plant to microbe. A majority of these SNPs were synonymous. However, some were non-synonymous, including the histidine decarboxylase SNP (B2R_16511) mapping to both chromosomes 6 and 11 (Figure 7). *Streptomyces* SNPs that were more abundantly associated with the wild tomato allele on chromosome 4 included an antibiotic resistance gene (daunorubicin/doxorubicin, B2R_28992) and maltooligosyl trehalose synthase (B2R_07820) among others.

**Figure 7:**
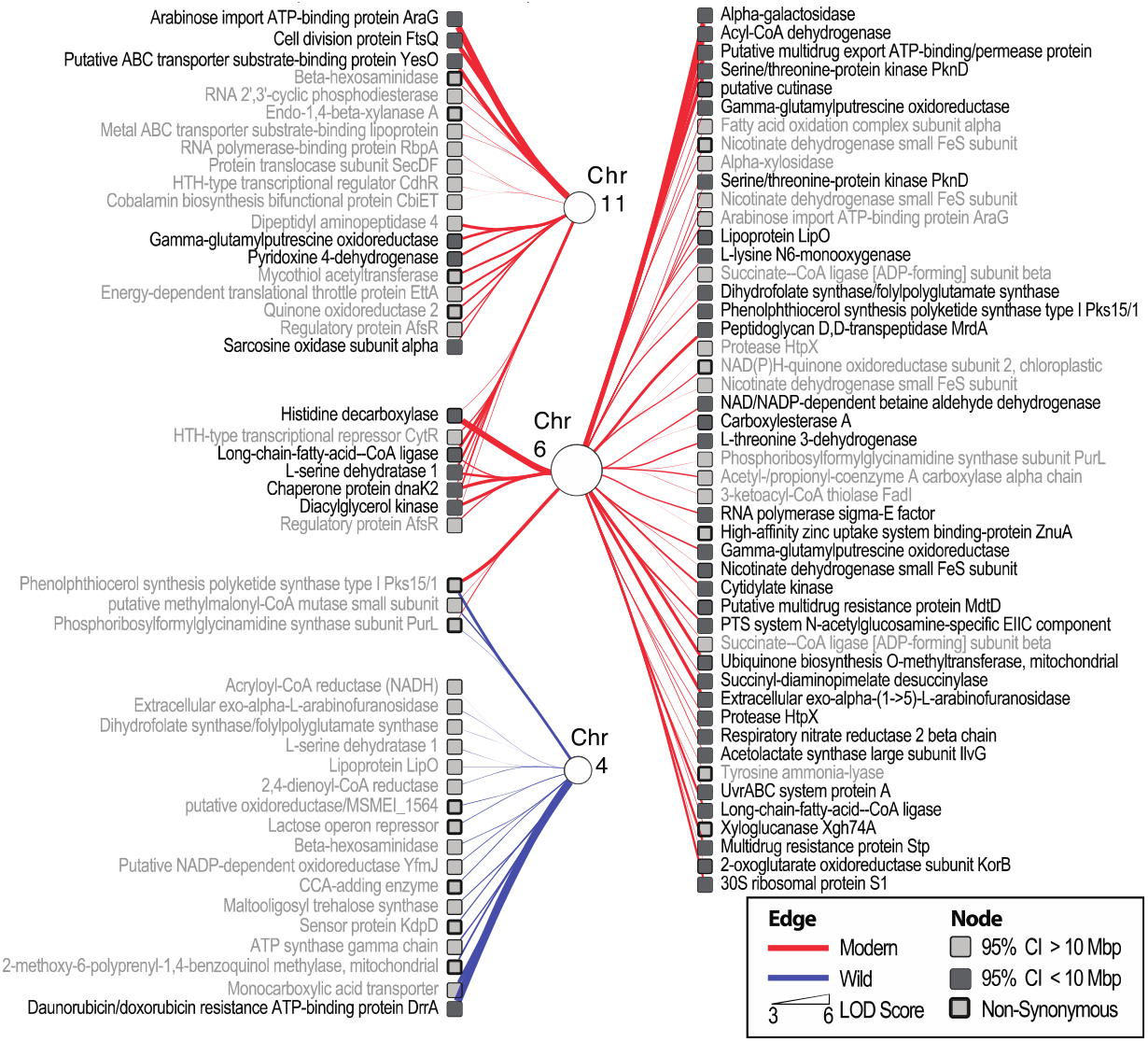
94 unique SNP within annotated genes on the Streptomyces contigs that mapped to QTL positions on tomato chromosomes 4, 6 and 11. The figure depicts various features of both the QTL analysis and the SNPs. In particular, the circular nodes represent the tomato chromosomes 4,6 and 11. The size of these nodes is relative to the number of outgoing edges, which represent individual QTLs. The width of edges is relative to the LOD score and are color coded depending on whether a QTL is linked to increased abundance in the modern or wild allele. The Streptomyces SNPs, which were the microbial molecular features mapped as QTLs, are represented by square nodes with annotations alongside. Those SNPs with confidence intervals <10 Mbp are shaded in dark. Non-synonymous SNPs have a thick border edge.

Similarly, we investigated the 324 *Cellvibrio* SNP QTLs within annotated genes whose mapping coincided with the previously identified *Cellvibrio* contig QTLs to chr. 1 and 9. Again, numerous SNV QTLs were identified in genes were related to sugar catabolism, including a gene encoding an extracellular exo-alpha-(1->5)-L-arabinofuranosidase (B2R_16093), fructose import FruK (B2R_22268), a cellulase/esterase-encoding *celE* homologue (B2R_11067), and genes involved in malate (B2R_18213), mannonate (B2R_14081), xyloglucan (B2R_10668) and xylulose (B2R_22179) metabolism. Furthermore, many additional SNP QTL were identified in genes related to vitamin and cofactor metabolism as well as sulfur and iron metabolism. In particular, these included genes for a phosphoadenosine phosphosulfate reductase (B2R_15720), vitamin B12 transporter BtuB (10 different genes, see Supplemental Table 17), a siroheme synthase (B2R_24033), a pyridoxal phosphate homeostasis protein (B2R_17481), a heme chaperone HemW (B2R_12751), a hemin transport system permease protein HmuU (B2R_09175), a Fe(2+) transporter FeoB (B2R_19968), a biotin synthase (B2R_30007), a catecholate siderophore receptor Fiu (B2R_17486), and a Fe(3+) dicitrate transport ATP-binding protein Fec (B2R_09176) (Supplemental Table 17). Taken together, this analysis suggests that a shotgun metagenomic approach integrated with quantitative plant genetics can be instrumental in a high-throughput manner to discover the reciprocal genetic links between plant and microbial metabolisms, such as those identified here for polysaccharides, trehalose, iron, vitamin, amino acid, and polyamine metabolism.

## 3. Discussion

Breeding for microbiome-assisted crops is a complex phenomenon encompassing ecological, evolutionary, and cultural processes. What constitutes a desirable trait for selection is context-dependent and differs between societies, crops and locations^61^. As society grapples with modern challenges such as a rapidly changing environment, water scarcity and land degradation, it is becoming increasingly clear that a new era of trait selection is needed with increased focus on sustainability and microbiome interactions^62–65^. In this regard, it is also time to reckon with the consequences of historic yield-centric trait selection and accompanying genomic sweeps^31^, especially with regards to plant-microbe interactions. Current approaches to investigating the genomic architecture determining microbiome assembly rely primarily on mutational studies in known genes and pathways. More recently, studies leveraging the natural variation within plant populations have been used to conduct GWA and QTL of the leaf^66,20^ and rhizosphere^18^. To date, the microbiome has been primarily characterized through amplicon sequencing, thereby providing limited functional resolution of microbiome structure. Increasing the resolution of phenotyping of quantitative traits has been shown to improve the precision and detection of QTLs^67^. Thus, integrating microbial genomics into microbiome QTL analysis plays dual purpose; increasing the ecological resolution with which microbial traits may be mapped, and second, affording the identification of the reciprocal microbial adaptations that drive plant-microbe interactions. In this investigation, we addressed these challenges by integrating amplicon and shotgun metagenome sequencing to identify microbiome QTLs.

One major difference between the amplicon and contig QTL analysis is the number of lineages for which QTLs were identified. In this regard, amplicon-based sequencing provided a broader picture and was able to capture QTLs of both abundant and relatively rare rhizobacterial lineages. In contrast, the majority of contig QTLs mapped to the most predominant lineages, yet failed to identify QTLs for more rare lineages. Nevertheless, besides the fact that the shotgun-based approach provided functional insights into the associated bacterial taxa, the size of the 95% confidence interval of the QTL region was significantly smaller using contig QTLs, with a median size of just 6.47 Mbp compared to 58.56 Mbp for the amplicon-based QTL regions. Furthermore, for *Streptomyces*, the number of unique QTLs identified was greater in the contig-based approach. Thus, we identified a trade-off between amplicon and shotgun-based technologies, whereby amplicon sequencing provides a deeper view into broad community structure, whereas shotgun-based approaches provided a more nuanced picture. In particular, the smaller regions identified by our contig-based metagenome mapping provided considerably more functional insights as it enabled us to analyze the genomic content contained in the regions linked to *Cellvibrio* and *Streptomyces*.

The increased QTL mapping resolution provided by shotgun-based phenotyping of the microbiome combined with SNP analysis provided a novel approach to leverage both the host diversity of the RIL and the natural microbiome population diversity to disentangle the reciprocal genomic adaptions between plants and natural microbiomes. For example, understanding the driving forces driving the abundances of rhizospheric *Streptomyces* is of increasing interest and has been linked to both iron^68^ and water limitations^53^. Here, we pinpointed the genetic basis for these interactions among the short list of highly expressed root-specific tomato genes linked positively to *Streptomyces* abundance including both aquaporin and FIT. More specifically, the aquaporin (SlTIP2.3) has the highest fold change in the roots of all tonoplast intrinsic proteins in the tomato genome^32,33^, while the FIT gene has been shown to largely control iron homeostasis in tomato^34,35^.

In addition to these high priority genes, many other key genes were identified in these regions. Those previously shown to contribute to microbiome assembly included 1-aminocyclopropane-1-carboxylate oxidase, which plays a central role in plant regulation of various processes including bacterial colonization and root elongation^69^ and alpha-humulene/(-)-(E)-beta-caryophyllene synthase, a terpene known to modify microbiome structure^39^. In addition, numerous genes related to growth, development and cell wall loosening^70^ known to be involved in microbial colonization^71^ and aluminum-activated malate transporter, which has been linked to microbiome mediated abiotic stress tolerance^40^.

The historic impact of domestication on genomic regions linked to microbiome assembly is also apparent (Figure 6, Supplemental Table 14, and Supplemental Figure 1). However, the processes and consequences of these sweeps, and possible subsequent recombination events on microbiome assembly remain unclear. In particular, the discontinuity of sweeps in microbiome QTL regions suggests that evolutionary pressure for recombination of key (microbiome associated) traits, such as iron homeostasis and water transport, may have acted against selective sweeps. The results obtained here provide the means to illuminate such complex eco-evolutionary questions, forming the basis of integrating the microbiome into the classic genotype by environment model of host phenotype^10^.

From the microbial perspective, the increased resolution in QTL analysis afforded by our shotgun-based approach also provided a window into the host-specific bacterial adaptations to wild and modern alleles. In particular, the SNP QTL analysis demonstrated that genes related to the degradation of various plant-associated polysaccharides in *Streptomyces* were highly associated with various modern tomato alleles. It should be further noted, that many other functions were identified in both plant and microbe, such as trehalose metabolism, polyamine metabolism and acetolactate synthase, suggesting either a direct link through cross-feeding^72^ or signaling^73^, or perhaps shared ecological pressures. While the microbial adaptations related to polysaccharides^74^, vitamins^75^ and iron metabolism^46,68^ are well documented in relation to plant colonization, here we demonstrate for the first time that the reciprocal adaptations that drive plant-microbe interactions can be investigated simultaneously to uncover their genetic architecture in both host and microbiome.

## 4. Methods

### 4.1 Greenhouse and Lab work

#### 4.1.1 Recombinant inbred line population

100 lines of an F8 recombinant inbred line (RIL) population derived from the parental lines *Solanum lycopersicum* cv. Moneymaker (Modern) and *Solanum pimpinellifolium* L. accession CGN14498 (Wild) were used^23^. A high density map produced from this population was used to map QTLs^26^.

#### 4.1.2 Growth conditions for RIL

The soil was collected in June 2017 from a tomato greenhouse in South-Holland, The Netherlands (51°57’47”N 4°12’16”E). The soil was sieved, air dried, and stored at room temperature until use in 2019. Before the beginning of the experiment, soil moisture was adjusted to 20% water by volume using deionized water. All soil was homogenized by thorough mixing and allowed to sit, covered by a breathable cloth, in the greenhouse for one week prior to potting. The soil was then homogenized once again and then potted. Each pot was weighed to ensure all pots were 175g±0.5 (wet weight). Duplicate pots for each accession were planted, as well as 6 replicates of each modern and wild parental accession, and 8 bulk soil pots that were left unseeded. Each replicate was prepared simultaneously. Planting was done separately representing biological replicates.

In each pot, 3 seeds were planted in a triangular pattern to ensure the germination success for all pots. The first seedling to emerge in each pot was retained and others were removed after germination. All pots were randomly distributed in trays containing approximately 10 plants. Throughout growth, careful attention was given to randomize the distribution of plants. First, tray location and orientation with relation to each other were randomized on a nearly daily basis. In addition, the distribution of plants within trays was randomized three times during growth. All pots were kept covered until germination, which was scored daily. After germination, plants were visually monitored and watered at the same rates. To minimize the impact of environmental differences between pots on microbiome composition, the watering regime for all plants was standardized and leaks from the bottom of the pot and overflows were completely prevented. To achieve this, a minimal volume (2.5 mL to 5.0 mL) of water was used at each watering. This strategy was successful as washout was never observed. Moisture content was measured by weighing the pots at the middle and end of the experiment to ensure all pots had similar moisture contents.

#### 4.1.3 Harvesting and processing of plant materials

All plants had between 5-7 true leaves at harvest (Supplemental Table 1). Plants were gently removed from the pot and roots and were vigorously shaken. Soil that remained attached to the roots after this stage was considered the rhizosphere. The remaining bulk soil and rhizosphere (plus roots) fractions were weighed. The root and attached rhizosphere fraction were treated with 4 mL of lifeguard, vortexed and sonicated. Roots were then removed. The remaining rhizosphere sample was then stored in LifeGuard Soil Preservation Solution (Qiagen) at -20 °C until DNA extraction.

The dry weight of shoots was measured after drying at 60°C. The dry weight of the bulk soil was measured after storing at room temperature in open paper bags for 1 month. The DNA was extracted using the DNeasy PowerSoil extraction kit (Qiagen). The protocol was optimized for the soil in the following manner: each sample was vortexed and then a volume of approximately 1.5 mL was transferred into 2 mL tubes. This subsample was centrifuged at 10,000g for 30 seconds such that a pellet was formed. The supernatant was removed, and a new subsample was transferred, and centrifuged until the total volume of the original sample, without sand, had been transferred to the 2 mL tubes. The resulting pellet was recalcitrant to disruption through bead beating, and therefore was physically disrupted by a pipette tip before proceeding with DNA extraction protocol. In test samples, DNA extractions from the sand fraction yielding no, or marginal levels of DNA.

### 4.2 Amplicon and shotgun metagenomics analysis

#### 4.2.1 rRNA amplicon sequence processing

All DNA was sent to BaseClear (Leiden, The Netherlands) for both 16S and 18S 300 bp paired-end amplicon sequencing (MiSeq platform). MiSeq primers targeted the V3-V4 region of Bacteria: 341F CCTACGGGNGGCWGCAG, 805R GACTACHVGGGTATCTAATCC. In total, 20,542,135 16S read pairs over 225 samples were generated. The raw reads were processed using the DADA2 workflow (v1.14.1) to produce amplicon sequence variants (ASV) and to assign taxonomy^76^. ASVs tagged as non-bacterial, chloroplast, or mitochondria were removed. Next, ASV counts were normalized using the cumulative sum scaling (CSS) and filtered based on the effective sample size using the metagenomeSeq package (v1.28.2)^77^. Differential abundances between rhizosphere and bulk soil were determined using the eBayes function from the limma package. Enriched rhizosphere ASVs with a greater than log(2) fold change in abundance were analyzed based on their presence and absence, standard deviation and mean values. Using these statistics, stochastic ASVs (<50% of samples) were removed from further analysis.

#### 4.2.2 Metagenomics analysis

For the one set of replicates for each accession, paired-end sequence read libraries were generated in the length of 150 bp per read on NovaSeq paired-end platform by BaseClear B.V. Demultiplexing was performed before the following analysis. It is computationally expensive to assemble the 114 read libraries all at once. Therefore, a strategy of (merging) partial assemblies was undertaken. Two assemblers were used to create the assembled contigs, namely SPAdes (version 3.13.2)^78^ and MEGAHIT (version 1.2.9)^79^. Assembly quality was assessed by running MultiQC (version 1.8)^80^ with Quast Module^81^(Supplemental Figure 2). First, 6 modern parents, 5 wild parents and 1 bulk soil sample were co-assembled via SPAdes with the metagenomic mode and parameter of -k 21,33,55,99, generating the first assembly (A1). Subsequently, a second assembly (A2) was done using the unmapped reads from the remaining metagenomes using MEGAHIT with the parameter of --k-list 27,33,55,77,99. The third assembly (A3) was performed similarly as A2, however included the unmapped reads, ambiguously mapped reads, and mapped reads with a low mapping quality score (MapQ < 20) (Supplemental Table 18). Read mapping was done with BWA-MEM with default settings^82^ and SAMtools was used to convert resulting SAM files into sorted and indexed BAM files (version 1.10). Extraction of these reads were conducted by samtools bam2fq. Redundancy between assemblies was evaluated by alignment to A1 via nucmer package of MUMmer with --maxmatch option (version:4.0.0)^83^.

Firstly, 111.5 Gbp of reads from the parental samples were assembled, labelled as A1 and yielded a total assembly length of 8.6 Gbp with the largest contig of 933.0 kilobase pairs (Kbp). After aligning the reads from RIL samples to A1, unmapped reads, ambiguously mapped reads, and mapped reads with a low mapping quality score (MapQ < 20) were retrieved and assembled, yielding the second and third assembly (A2 and A3). Specifically, A2 stemmed from solely the unmapped reads while A3 included the ambiguously mapped reads and mapped reads with MapQ < 20 in addition to the unmapped reads. A2 and A3 produced a total assembly length of 9.6 Gbp and 14.0 Gbp, with the largest contig of 56.2 Kbp and 86.3 Kbp respectively. There were 1.2, 2.0 and 2.8 million contigs with the length over 1 Kb for A1, A2 and A3 respectively. In particular, 912 contigs in A1 were greater or equal to 50 Kbp whereas 1 or 2 such large contigs were successfully assembled in A2 or A3. The detailed assembly statistics is given in Supplemental Table 18 and the numbers of contigs with different ranges of length for each assembly are presented in Supplemental Figure 2.

The sequence similarities of the contigs in each assembly (≥ 1 Kbp) were compared using the nucmer package in MUMer. No contigs in A2 were reported to share an overlapped region with A1, therefore contigs in A1 and A2 could be merged directly. When A3 was aligned to A1, 1.1% of the total length (≥ 1 Kbp) of A3 was reported to be overlapped with A1, however, only 18 contigs from A3 were 100% identical to regions in larger contigs in A1. The sensitivity of filtering the overlapping contigs was evaluated by a benchmarking test using a random RIL sample to calculate the mapping rates (Supplemental Figure 3). 83.4% reads were mapped to A1+A3 at MapQ ≥ 20 without filtering. Excluding the contigs from A3 that were completely and identically covered by A1, the mapping rate was nearly the same as the one without filtering. Nevertheless, the removal of all aligned contigs in A3 resulted in a slight drop of mapping rate to 82.6%. To conclude, the final assembly was determined as A1+A3 with the 18 redundant contigs from A3 removed.

To assess the overall assembly quality and quantify the abundance of contigs among all samples, metagenomic reads were mapped to A1, A1+A2 and A1+A3 (deduplicated) respectively. Afterwards, the mapping rates were calculated for the mapped reads with MapQ > 20 in each sample. As shown in Supplemental Figure 4, approximately 70% reads among rhizosphere samples could be mapped to A1, while the mapping rates were 55% to 65% in the bulk soil samples. With the unmapped reads assembled and added to A1, the mapping rates for A1+A2 increased by 10%. The read recruitment was further improved by assembling and adding ambiguously mapped reads and mapped reads with low MapQ in the final assembly (A1+A3). A1 as well as de-replicated A3 were merged to acquire the final assembly. All the ‘contigs’ mentioned below are referring to the contigs in this final assembly.

#### 4.2.3 Binning of metagenomic contigs

Metabat2 (version 2:2.15)^84^ was used for assigning the contigs into genomic bins. Based on tetra-nucleotide frequency and abundance scores, 588 genomic genomics bins were generated. Afterwards, genomic quality of those genomes was evaluated by CheckM (version: 1.1.1)^28^ with the command “checkm linage_wf” (Supplemental Table 9). The 33 genomes displaying the completeness larger than 90% and contamination smaller than 5% were used for further study as quantitative traits.

#### 4.2.4 Making phenotype files based on contig depth

Read counts for each position on the assembled contigs were acquired using bedtools genomecov (version: 2.29.2)^85^. A custom Python script was applied to calculate the average depth (defined as the number of total mapped reads divided by contig length) and coverage (defined as the number of covered base pairs divided by contig length) of every contig. Furthermore, the average abundance of contigs assigned into a bin was calculated for the high-quality genomic bins detected by CheckM^28^.

#### 4.2.5 Feature selection

Average depths of the contigs were first normalized using the cumulative sum scaling (CSS) and filtered based on the effective sample size using metagenomeSeq package (v1.28.2)^77^. Differential abundance analysis was performed by moderated t-tests between groups using the makeContrasts and eBayes commands retrieved from the R package Limma (v.3.22.7)^86^. Obtained P-values were adjusted using the Benjamini–Hochberg correction method. Differences in the abundance of contigs between groups were considered significant when adjusted P-values were lower than 0.01 (Supplemental Table 19).

In either comparison, the contigs that were significantly enriched in the rhizosphere were gathered and regarded as the statistically rhizosphere-enriched contigs after removing the replicated ones. To perform QTL analysis for the abundance of these enriched rhizosphere contigs, only the contigs with biological meanings were kept, i.e. the log (2) fold-change of mean values for the normalized abundances of RIL and bulk samples should be greater than 2, and the contig should be in enough depth with at least the mean value of a group larger than 1. This selection step resulted in 1249 rhizosphere-enriched contigs in the end. The statistics of the filtered normalized abundance were further inspected based on the presence and absence of contigs, standard deviation and mean values of the counts.

#### 4.2.6 Taxonomic and functional annotation of the metagenome

Taxonomic classifications were assigned to the contigs in the final assembly using Kraken2 (version: 2.0.8)^29^ based on exact k-mer matches. A custom Kraken2 database was built to contain *RefSeq* complete genomes/proteins of archaea, bacteria, viral, fungi and protozoa. Univec_Core was also included in the custom database (20200308). Using the Kraken2 standard output, a python script based on TaxonKit^87^ was utilized to add full taxonomic names to each contig in the format of tab-delimited table. 76.22% of the contigs > 1kb were classified. Among the contigs > 10kb, up to 99.44% contigs were classified. Prokaryotic microbial genes were predicted by Prodigal (version: 2.6.3)^88^ with metagenomics mode. 10,246,55 genes were predicted from contigs > 1kb (Supplemental_Table_8). Open reading frames (ORFs) on contigs >10kb were annotated by prokka (v1.14.5) and the *Streptomyces* ASV5 bin (MAG.72) was further annotated by DRAM (v1.2.0) integrating UniRef, Pfam, dbCAN and KEGG databases^89^.

#### 4.2.7 Single Nucleotide Variant analysis

To investigate strain level QTLs, we mapped Single Nucleotide Variants (SNV) identified using inStrain on the 1249 contig enriched genomes. A total of 555,382 and 535,432 SNPs were identified in the modern and wild parental metagenomes respectively. Of these, 162,299 and 142,349 SNPs were unique to each dataset respectively, as they either contained only reference alleles or did not exceed the inStrain SNP calling thresholds. For each unique SNP locus, coverage in the other dataset was determined using SAMtools depth after read filtering with settings comparable to inStrain, and was considered identical to the reference allele frequency. Including the unique SNPs, this resulted in a final set of 697,731 SNPs. To select SNPs that showed differential reference allele frequencies between MM and P, first the difference in reference allele frequency (MM – P) was calculated per SNP. From the distribution of all SNPs, the 95% confidence interval (CI) was determined to select the 5% (30,911) most different SNPs (Supplemental Figure 5). SNPs were further selected using a Fisher’s exact test based on the allele read count differences between MM and P. P-values were sorted, and a final selection of 15,026 differentially abundant SNPs distributed over 1,037 contigs was obtained using a Benjamini-Hochberg false discovery rate (FDR) correction of 0.01. SNV allele read counts were extracted from the RIL dataset using the pysam Python package after filtering with settings comparable to inStrain.

#### 4.2.8 Quantitative Trait Locus Analysis

The QTL analysis linking selected amplicon, contig, bin, and SNV features with plant loci was performed using the R package R/qtl2^25^. Pseudomarkers were added to the genetic map to increase resolution, with a step distance of 1 Mbp between the markers and pseudomarkers. Plant genome probabilities were calculated using the genetic map with pseudomarkers, plant loci cross data and error probability of 1E-4. Plant locus kinship matrix was calculated as proportion of shared alleles using conditional allele probabilities of all plant chromosomes, which were calculated from the plant genome probabilities. A genome scan using a single-QTL model using a linear mixed model was performed on the SNP allele read counts as phenotypes, plant genotype probabilities as input variables and as covariates the number of leaves, harvest day, rhizosphere soil weight (g), soil starting weight (g) and plant dry weight (g). The log_10_ likelihood (LOD) score was determined for each plant locus SNP allele combination. A permutation test was performed with 1000 permutations to assess the distribution of the LOD scores. The 95% quantile was used as threshold for the selection of LOD peaks, as well as a P = 0.95 Bayes credible interval probability.

### 4.3 Independent validation of QTLs through bulk segregant analysis

To validate the QTLs, 33 *Solanum lycopersicum* cv. Moneymaker (Modern), 30 *Solanum pimpinellifolium* L. accession CGN14498, and 77 RIL accessions (with replicates of 4 each) were grown and their microbiomes characterized through 16S sequencing. Parental lines and RIL accessions were germinated in pots filled with 300 g agricultural soil. For each accession, were planted with six plants per replicate pot. The plants were arranged randomly in the growth chamber (25°C, 16h daylight) and watered every day. Bulk soil samples without plants were used as controls (N = 31).

Rhizospheric soil was collected according to standard methods^90^. In order to synchronize the developmental stage, the plants were harvested after 21 days, or when the 3^rd^ trifoliate leaf was reached. The soil loosely attached to the roots was removed and the entire root system was transferred to a 15 mL tube containing 5 mL LifeGuard Soil Preservation Solution (MoBio Laboratories). The tubes were vigorously vortexed and sonicated. Subsequently, the roots were removed and at least 1 g (wet weight) of rhizospheric soil was recovered per sample for RNA extraction. For the bulk soil samples, approximately 1 g of soil was collected and mixed with 5 mL of LifeGuard solution.

To extract rhizospheric DNA, PowerSoil Total DNA/RNA Isolation Kit (MoBio Laboratories, Inc., USA) was used in accordance with manufacturer’s instruction. Rhizospheric DNA was obtained using RNA PoweSoil DNA Elution Accessory Kit (MoBio Laboratories, Inc. USA). The quantity and quality of the obtained DNA was checked by ND1000 spectrophotometer (NanoDrop Technologies, Wilmington, DE, USA) and Qubit 2.0 fluorometer (ThermoFisher Scientific, USA). DNA samples were stored at -20°C until further use.

The extracted samples were used for amplification and sequencing of the 16S rRNA, targeting the variable V3-V4 (Forward Primer= 5’CCTACGGGNGGCWGCAG-3’ Reverse Primer= 5’-GACTACHVGGGTATACTAATCC-3’) resulting in amplicons of approximately ∼460 bp. Dual indices and Illumina sequencing adapters using the Nextera XT Index Kit were attached to the V3– V4 amplicons. Subsequently, library quantification, normalization and pooling were performed and MiSeq v3 reagent kits were used to finally load the samples for MiSeq sequencing. For more info please refer to the guidelines of Illumina MiSeq System. The RDP extension to PANDASeq^91^, named Assembler^92^, was used to merge paired-end reads with a minimum overlap of 10 bp and at least a Phred score of 25. Primer sequences were removed from the per sample FASTQ files using Flexbar version 2.5^93^.

## Supporting information

Supplemental Tables and Figures Description

## 5. Data availability

The sequencing data generated in this study are available under ID BioProject ID PRJNA787039 (16S amplicons) and PRJNA789467 (shotgun metagenomics). Bacterial ASV reference sequences, and metagenome assembled genomes are available at https://osf.io/f45ek/.

## 6. Code availability

The code used in the analysis can be found at https://osf.io/f45ek/.

## 7. Author contributions and acknowledgements

The study was conceived and designed by BOO, VJC, WLi, MHM and JMR. The greenhouse experimentation and lab work were conducted by BOO, SSF, VC, VJC, AN. Contributions to data analysis came from BOO, TG, XP, EvdW, NS, AK, VC, VJC, BLS, MHM. The manuscript was drafted by BOO, BLS, MHM and JMR. All authors contributed to the revision and agreed upon the final draft. The project was financially supported, in part, by the NWO-TTW Perspective program BackToRoots (TTW-project 14218 to JMR, VJC, VC and BOO), by the NWO-Gravitation program MICRop (to JMR, MHM), a National Institutes of Health (NIH) Genome to Natural Products Network supplementary award (no. U01GM110706 to MHM), a ZonMW Enabling Technologies Hotel project (no. 40-43500-98-210 to MHM), a Senescyt fellowship awarded to SSF, and by internal funding from the Netherlands Institute of Ecology.

