## Supplementary figures and images for "Disentangling the genetic basis of rhizosphere microbiome assembly in tomato"

### Supplemental Figure 1

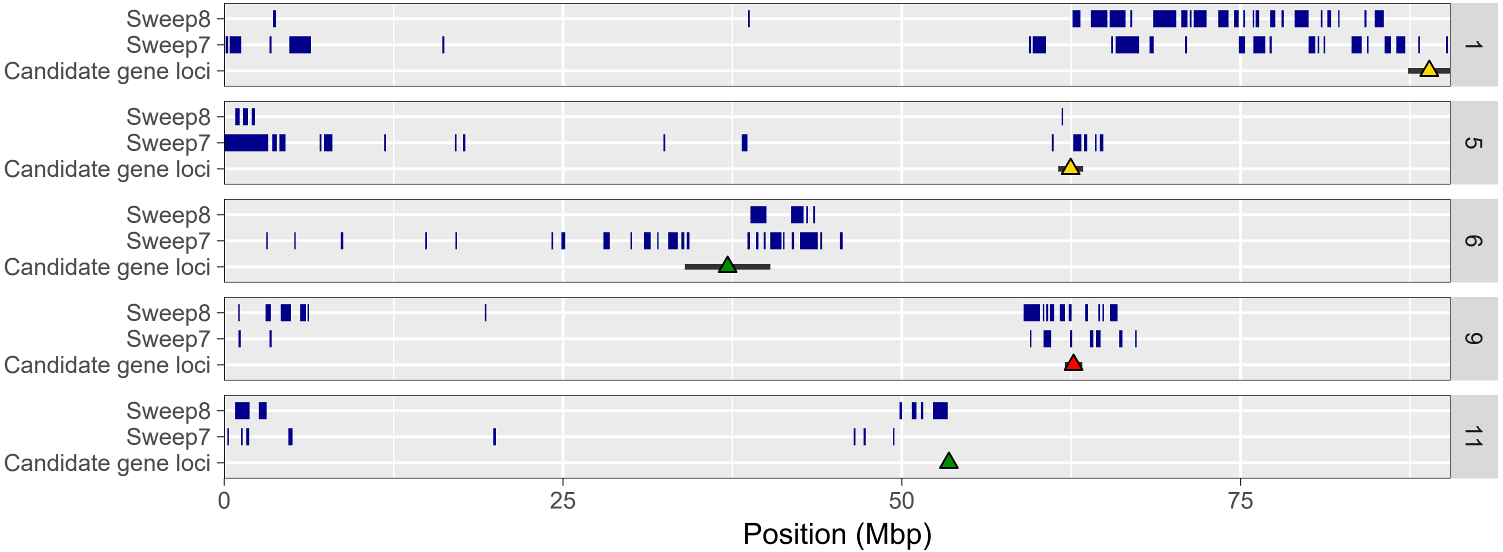
