## Supplemental_Figure_2-5 for "Disentangling the genetic basis of rhizosphere microbiome assembly in tomato"

### Slide 1
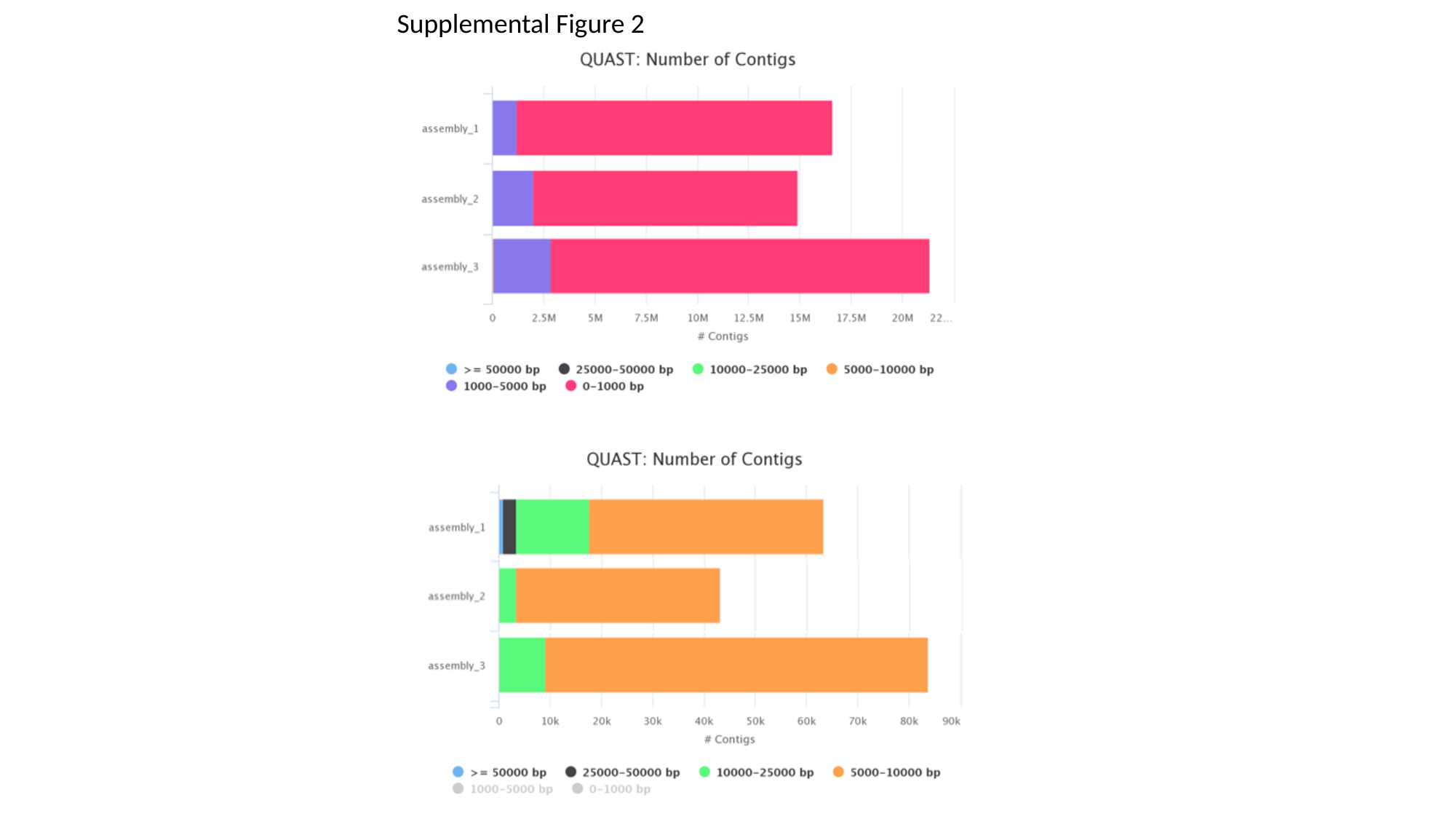

Supplemental Figure 2

### Slide 2
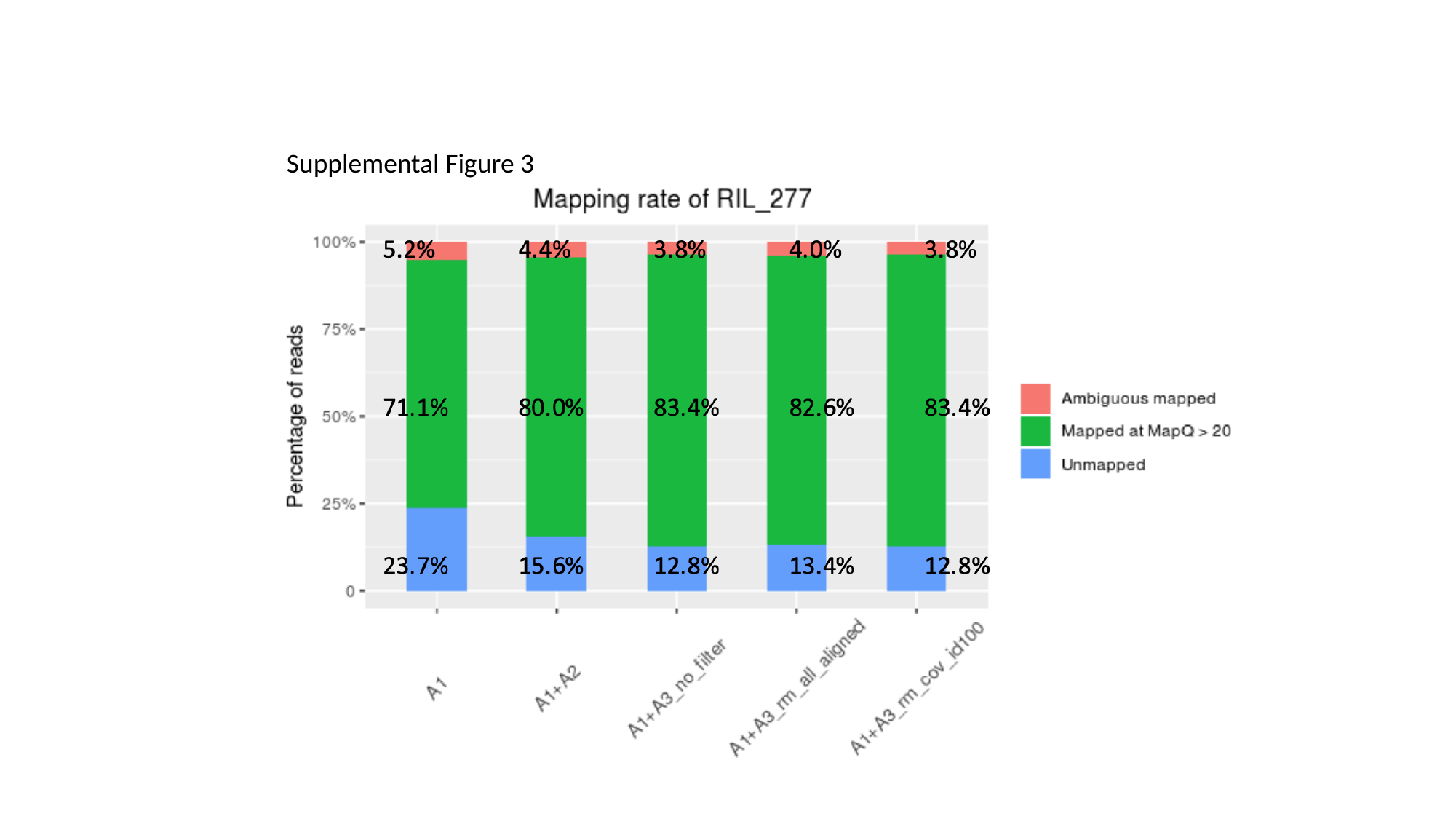

Supplemental Figure 3

### Slide 3
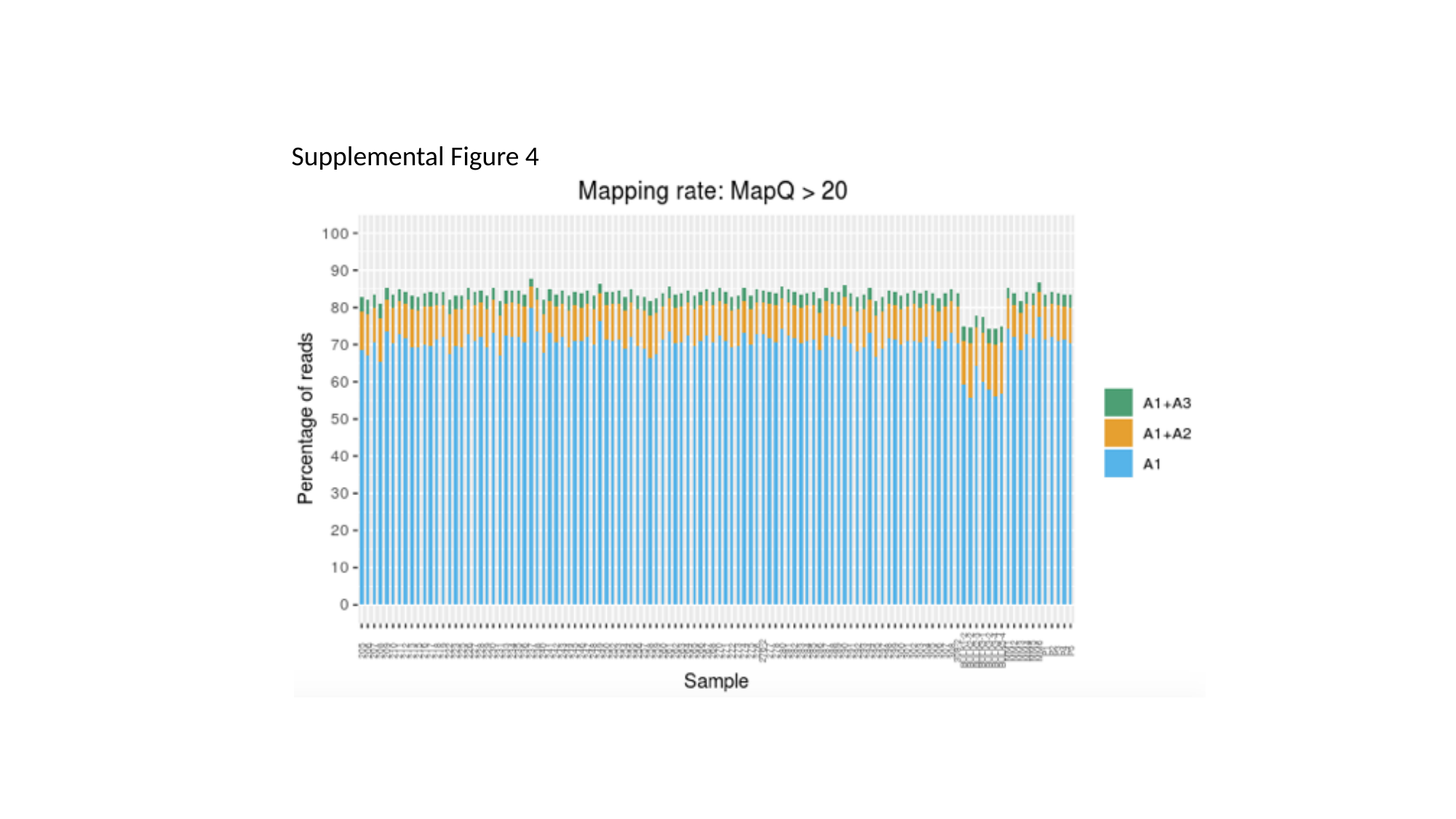

Supplemental Figure 4

### Slide 4
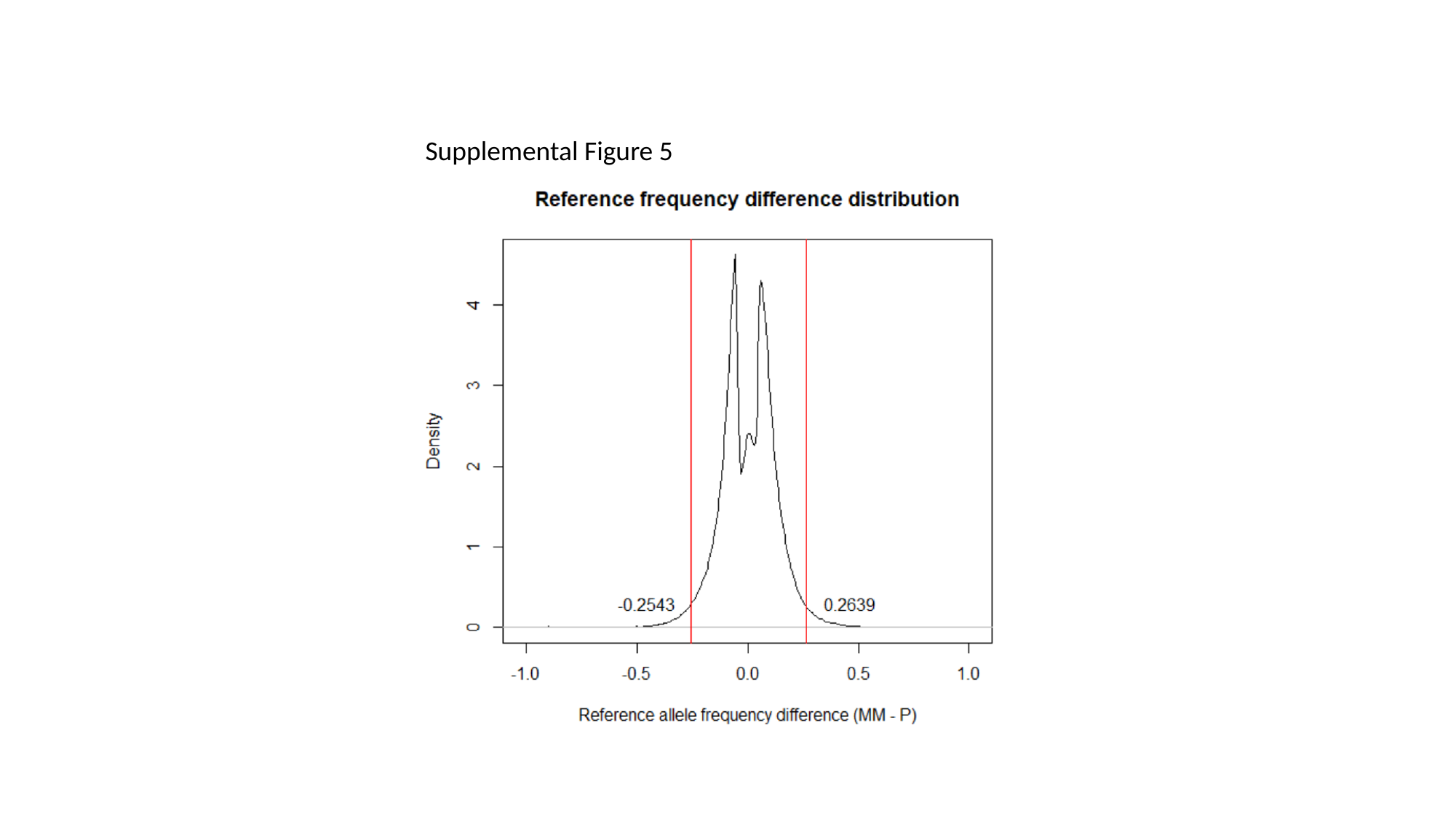

Supplemental Figure 5
