## Supplemental Tables and Figures Description for "Disentangling the genetic basis of rhizosphere microbiome assembly in tomato"

**Supplemental Tables**

**Metadata**

1. Supplemental_Table_1_RIL_metadata_replicates_5_19_2019.txt

Various meta data collected for each pot harvested. Includes the unique ID, the replicate (Color coded), the germination day and date, as well as the days of growth. Furthermore, this metadata sheet also includes the date of harvest number, the number of leaves at harvest (Leaves), the mass of the rhizosphere harvested in grams (Rhizosphere), the mass of the bulk soil remaining in grams (Soil_Before), and the plant dry weight in grams (PDW).

**Amplicon sequencing**

2. Supplemental_Table_2_asv_raw_and_taxonomy.csv

The metadata on each ASV including its raw abundance in each sample. The first columns are the kingdom, phylum, class, order, family, genus, species. The remaining columns represent the sample IDs.

3. Supplemental_Table_3_asv_css.csv

The abundance of ASVs after CSS normalization and filtering with each columns representing a different sample.

4. Supplemental_Table_4_asv_css_statistics_rhizosphere.csv

Some basic statistics about each ASV, including the number of zeros within the RIL population (zeroes), the number of non-zero abundances (nz), the mean (mean), the mean when excluding zero counts (nz_mean), the sum, minimum (min), maximum (max), the normalized standard deviation (norm_sd), the normalized standard deviation of non-zero counts (norm_sd_nz) and finally the a classification into either core, flexible or uncommon (ecology).

5. Supplemental_Table_5_sequence_table_nochimeras.fna

A fasta file for the ASVs identified in samples harvested on 5/19/2019.

6. Supplemental_Table_6_found_peaks_r.txt

The ASV QTLs identified including the ASV, the chromosome (chr) and position in Mbps (pos), the LOD score (lod), the low (ci_lo) and high (ci_hi) 95% confidence intervals in Mbps, the effect, heritability, whether a wild or modern allele was observed at that position. In addition, taxonomical information is included.

**Shotgun metagenomics**

7. Supplemental_Table_7_contig_10kb_taxonomy.tsv

The taxonomy output from Kraken on the contigs 10kb and greater in size.

8. Supplemental_Table_8_new_checkm_lineage_wf.tsv

The taxonomy output and quality assessments of CheckM of the metagenome assembled bins

9. Supplemental_Table_9_found_peaks_for_bin_permu1000_full.tsv

The bin QTLs identified including the bin id, the chromosome (chr) and position in Mbps (pos), the LOD score (lod), the low (ci_lo) and high (ci_hi) 95% confidence intervals in Mbps, the effect, heritability, whether a wild or modern allele was observed at that position.

10. Supplemental_Table_10_peak_contig_for_report.tsv

The contig QTLs identified including the contig id, the chromosome (chr) and position in Mbps (pos), the LOD score (lod), the low (ci_lo) and high (ci_hi) 95% confidence intervals in Mbps, the effect, heritability, whether a wild or modern allele was observed at that position. In addition, taxonomical information is included.

**Bulk segregant analysis**

11. Supplemental_Table_11_ BSA_RIL1_summary.xlsx

A table summarizing the output from the independent population of RIL accessions. The metadata includes the sample unique identifier (Replicate), whether a sample was a bulk soil, parental (*Solanum lycopersicum* cv Moneymaker, M; *Solanum pimpinellifolium*, P), the RIL accession number (three digit code), or bulk soil (Bulk). In the next three columns, the SNP information for markers 2274 linked to *Streptomyces*, and markers 464 and 3142 linked to *Cellvibrio* (A allele – *Solanum lycopersicum* cv Moneymaker, B allele - *Solanum pimpinellifolium*). Finally, the last two columns are the *Streptomyces* (ASV3) and *Cellvibrio* (ASV9) CSS normalized abundances. The ASV numbering is different as this was an independent experiment.

**Tomato genome analysis**

12. Supplemental_Table_12_Candidate_genes_microbeQTLs_tomato_apr2021_final_plus_sweeps.xlsx

A summary of the five prioritized QTL regions. In the first sheet ‘Info’, key information about each QTL is summarized including a unique ID (QTL_ID), the left and right side of the confidence interval as well as its size, genus and number (and type) of combined QTLs to form this prioritized region are provided. In subsequent sheets, the positions, annotations, expression patterns^26^ and gene sweeps are provided^28^.

13. Supplemental_Table_13_RTR_Analysis.xlsx

A summary of the five prioritized QTL regions in a single sheet and with additional calculations including the log_2_ of the leaf/root ratio. An arbitrarily small value (0.1) was added to both root and shoot calculations to prevent fold calculations approaching negative infinity.

**Annotations**

14. Supplemental_Table_14_476_contigQTLs_DBCAN_output.tsv

Focused annotations for the contigs with QTLs including, HMMER, Hotpep, DIAMOND as well as a unique gene ID.

15. Supplemental_Table_15_microQTL_contig_ORF_table_with_taxonomy.txt

Annotations for all contigs that were 10 kb and greater.

16. Supplemental_Table_16_contigs_QTL_antiSMASH_summary_061221.numbers

Summary of output from AntiSmash.

**SNV analysis**

17. Supplemental_Table_17_focused_SNV_QTL_found_peaks_annotations_210708_newORF.xlsx

The SNV QTLs identified including the SNV id, the chromosome (chr) and position in Mbps (pos), the LOD score (lod), the low (ci_lo) and high (ci_hi) 95% confidence intervals in Mbps, the effect, heritability, whether a wild or modern allele was observed at that position. In addition, taxonomical information is included, as well as annotation.

**Assembly**

18. Supplemental_Table_18_Assembly_statistics.xlsx

Assembly statistics from the three assemblies and their merged results.

19. Supplemental_Table_19_Contig_Feature_Selection.xlsx

Determining which contig features were selected

**Supplemental Figures**

1. **Supplemental Figure 1 –** Overlay of gene sweeps on QTL positions

The distribution of gene sweeps linked to the initial domestication and improvement for fruit (quality) traits (sweep7 and sweep8 respectively)^31^ overlaid with the prioritized QTLs on chromosomes 1, 5, 6, 9, 11.

2. **Supplemental_Figure_2** MultiQC Contigs across different assembly strategies

Bar plots generated by MultiQC showing the number of contigs with different ranges of length in the metagenomic assemblies. Figure S1A provides an overview of all the contigs and large contigs are focused in Figure S1B. The bars are color coded based on the length of contigs. “assembly_1”, “assembly_2” and “assembly_3” indicated the first, second and third assembly respectively. The third assembly yielded the greatest total number of contigs but most large contigs (≥ 10 Kbp) were successfully assembled in the first assembly.

3. **Supplemental_Figure_3** – Back-mapping of randomly sampled RIL

Mapping rates of RIL 277 in the benchmarking test on the filtering sensitivity of overlapping contigs. A1: the first assembly using reads from 11 parental (6 modern, 5 wild) and 1 bulk-soil samples; A2: The reads from the RIL metagenomes were mapped to assembly A1, and all unmapped reads were assembled; A3: Again, as with assembly A2, the third assembly used unmapped reads, but also included ambiguously mapped and low-quality mapped (MapQ < 20) reads from RIL samples. A1 and A2 were merged directly because there were no overlapping contigs, which was represented by “A1+A2” in the figure. The filtering of overlapping contigs in A3 was divided to 3 levels of stringencies: removing all aligned contigs (the most stringent), removing the overlapping contigs with 100% identity and coverage, and keeping all the aligned contigs (the loosest), which were represented by “A1+A3_rm_all_aligned”, “A1+A3_rm_cov_id100” and “A1+A3_no_filter” respectively in the figure.

4. **Supplemental_Figure_4** – Back-mapping of all samples

The mapping rates for three metagenomic assemblies. The first assembly and two merged assemblies were indexed and treated as the reference respectively in the backmapping for the metagenomic reads. For each sample, the number of reads with a mapping quality equal or greater than 20 were counted by using SAMtools and divided by the total number of reads per sample (including both reverse and forward reads). This figure shows that compared to the first assembly, the read recruitment for the merged assemblies were improved by adding unmapped reads, ambiguously mapped reads, and mapped reads with a low mapping quality score (MapQ < 20) from the RIL accessions. The mapping rates for the final assembly (A1+A3) were from 75% to 88%.

5. **Supplemental Figure 5 –** SNV feature selection

Distribution of difference in SNP reference allele frequency between the MM and P metagenomes (MM – P). Red lines indicate the 95% CI and the corresponding values indicate the significance thresholds. The 30,932 SNPs with reference allele frequency difference outside the 95% CI are used for further selection. The two peaks just around 0 arise from the addition of the SNPs that are called by inStrain in one dataset, but not in the other, and are thus assumed to comprise 100% reference alleles. This often leads to SNPs being recognized in one dataset with at most 95% reference allele frequency (5% SNP frequency is minimum requirement), while in the other dataset there is 100% SNP identity, resulting in the large peaks of barely different SNP loci. The SNPs in between are a result of identical reference allele frequencies, but this need not indicate similar variant alleles.
